# Genotype-specific transcriptional responses overshadow salinity effects in a marine diatom sampled along the Baltic Sea salinity cline

**DOI:** 10.1101/2021.11.04.467364

**Authors:** Eveline Pinseel, Teofil Nakov, Koen Van den Berge, Kala M. Downey, Kathryn J. Judy, Olga Kourtchenko, Anke Kremp, Elizabeth C. Ruck, Conny Sjöqvist, Mats Töpel, Anna Godhe, Andrew J. Alverson

## Abstract

The salinity gradient separating marine and freshwater environments represents a major ecological divide for microbiota, yet the mechanisms by which marine microbes have adapted to and ultimately diversified in freshwater environments are poorly understood. Here, we take advantage of a natural evolutionary experiment: the colonization of the brackish Baltic Sea by the ancestrally marine diatom *Skeletonema marinoi*. To understand how diatoms respond to low salinity, we characterized transcriptomic responses of *S. marinoi* grown in a common garden. Our experiment included eight genotypes from source populations spanning the Baltic Sea salinity cline. Changes in gene expression revealed a shared response to salinity across genotypes, where low salinities induced profound changes in cellular metabolism, including upregulation of carbon fixation and storage compound biosynthesis, and increased nutrient demand and oxidative stress. Nevertheless, the genotype effect overshadowed the salinity effect, as genotypes differed significantly in their response, both in the magnitude and direction of gene expression. Intraspecific differences included regulation of transcription and translation, nitrogen metabolism, cell signaling, and aerobic respiration. The high degree of intraspecific variation in gene expression observed here highlights an important but often overlooked source of biological variation associated with how diatoms respond and adapt to environmental change.

## INTRODUCTION

The salinity gradient separating marine and freshwater environments represents one of the major ecological divides structuring microbial diversity [1]. Differences in osmotic pressure impede marine– freshwater transitions, and as a consequence, transitions are generally rare, occur on longer evolutionary timescales [2, 3], and have repeatedly led to bursts of diversification in freshwater environments [4]. Comprehending the processes behind marine–freshwater habitat transitions is fundamental to both our understanding of lineage diversification and habitat structuring on evolutionary time-scales [5], as well as short term adaptive potential to climate change. The latter is rapidly altering marine environments: melting ice caps, altered precipitation patterns, and changes in oceanic currents have led to freshening of large regions as well as local changes in the seasonal or annual cycling of salinity regimes [6, 7]. Yet, little is known about the cellular processes underlying acclimation and adaptation to low salinity environments. Permanent establishment of ancestrally marine organisms in freshwaters depends on the ability of the individual colonists to survive the initial hypo-osmotic stress, acclimate to low salinity, and ultimately adapt to their new environment [8]. Consequently, such transitions are not likely to occur instantly, but are rather thought to happen gradually [4, 5]. Euryhaline species that are able to tolerate a wide range of salinities, or those inhabiting brackish environments, are most likely to succeed at marine–freshwater transitions. Thus, to understand the cellular processes behind marine–freshwater transitions, studies focusing on taxa that are most likely to colonize freshwater habitats are crucial.

Here, we take advantage of a natural evolutionary experiment currently underway: the colonization of one of the world’s largest brackish water bodies, the Baltic Sea, by the ancestrally marine diatom *Skeletonema marinoi* (Fig. 1A). Geologically, the Baltic Sea is young, with sea ice from the last glacial maximum having fully receded only 10 000 years ago and inundation of saline waters from the adjacent North Sea occurring just 8 000 years ago [9]. Today, freshwater input from rivers and precipitation, in combination with continued inflow of saline bottom-waters from the North Sea through the Danish straits, results in a latitudinal and vertical salinity gradient ranging from near fresh to fully marine conditions [9, 10] (Fig. 1A). This salinity gradient strongly structures biodiversity of aquatic biota on both the species and population levels [11–13], including diatoms [14, 15]. This includes *S. marinoi*, which is the dominant phytoplankton species and one of the main contributors to primary production in the area [16, 17]. Paleoecological evidence showed that *S. marinoi* has been present in the Baltic Sea since the marine inundation or shortly thereafter [18, 19]. Although *S. marinoi* is ancestrally marine [20, 21], it can tolerate a wide range of salinities and is common along the entire salinity gradient, from the North Sea coast to the upper reaches of the Baltic Sea [22]. Previous work showed reduced gene flow between a high-salinity North Sea population of *S. marinoi* and a low-salinity Baltic Sea population [22]. The Baltic population exhibited lower genetic diversity and optimal growth at lower salinity, consistent with local adaptation [22]. Thus, *S. marinoi* presents an excellent system for understanding the cellular mechanisms that govern adaptation of ancestrally marine diatoms to low salinity environments.

**Fig. 1.**
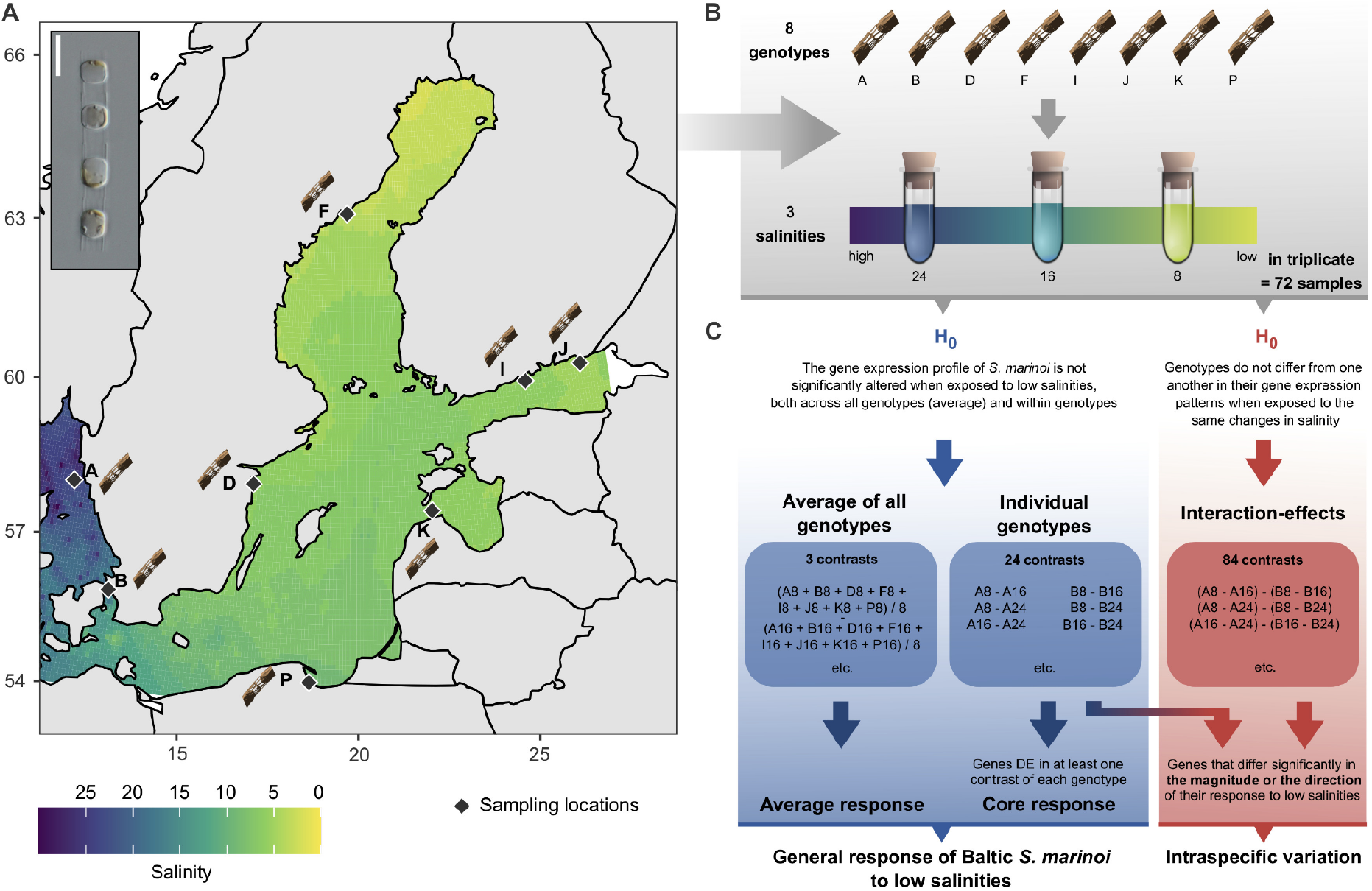
Experimental design. **a Field sampling.** Natural salinity gradient in the Baltic Sea based on salinity measurements from surface samples (0-10 m depth) and interpolated across the Baltic Sea for the period 1990-2020. Salinity measurements were downloaded from ICES (ICES Dataset on Ocean Hydrography, 2020. ICES, Copenhagen) and Sharkweb (https://sharkweb.smhi.se/hamta-data/). Diamonds identify sampling locations for *S. marinoi*. The inset figure shows a light micrograph of a *S. marinoi* culture (scale bar = 10 μm). **b Laboratory experiment.** Experimental design of the laboratory experiment carried out in this study. Eight strains of *S. marinoi* were exposed to three salinity treatments (8, 16 and 24) in triplicate, resulting in 72 RNA-seq libraries. **c Statistical analyses.** Overview of the null hypotheses and contrasts tested in this study. Our experimental design allowed characterization of the general response of *S. marinoi* to low salinities as well as intraspecific variation. The lower blue arrows indicate which data were incorporated in the average and core responses, which together were used to define the general response of *S. marinoi*. Genes with significant interaction effects were subdivided in two categories using logFC values of the genotype-specific effects (blue-red gradient arrow), distinguishing genes that differed significantly in either the magnitude or direction of their response to low salinities. The first category includes genes that were DE in one genotype but not the others, or that were DE in multiple genotypes but with significant differences in logFC values in the same direction. Genes of the second category were significantly upregulated in some genotypes, whereas they were significantly downregulated in other genotypes.

We combined a laboratory common garden experiment with RNA-sequencing (RNA-seq) to characterize the response of the euryhaline diatom *S. marinoi* to low salinity (Fig. 1). We collected eight genotypes of *S. marinoi* along the Baltic Sea salinity cline, exposed them to a range of salinities, and compared gene expression between high and low salinity treatments. Natural populations of *S. marinoi* exhibit a broad range of variability in ecophysiological traits [23], so the inclusion of multiple genotypes in our experiment allowed us to characterize potential variation in the salinity response as well, such as which aspects of the response are shared and which ones differ among genotypes.

## MATERIAL & METHODS

### Sample collection, culturing, and experimental design

We collected sediment samples from eight locations across the Baltic Sea (Fig. 1A) and stored them in the dark at 5 °C. We germinated *S. marinoi* resting cells into monoclonal cultures that were kept at their native salinity (Table 1) [24]. Cells were grown at 12 °C on a 12:12 light:dark light regime (30 μmol m^−2^ s^−1^ light intensity) for 12–26 months prior to the experiment. Strain identity was confirmed by sequencing the LSU d1–d2 rDNA gene (see Supplementary Methods for details).

**Table 1.** Details of the *S. marinoi* strains used in this study. The salinity values indicate the salinity of the natural sample from which the respective strains were isolated (‘original salinity’) and in which they were maintained prior to the experiment (‘culture salinity’). GPS coordinates indicated with an asterisk (*) represent approximate sampling locations.

| Collection ID | Strain ID | Field ID | Country | GPS (N/E) | Collector | Isolation date | Culture medium | Original salinity | Culture salinity |
| --- | --- | --- | --- | --- | --- | --- | --- | --- | --- |
| AJA304 | A.2.21b | A | Sweden | 58.02868/11.13738 (*) | A.Godhe | 2017-03-28 | L1 | 15-33 | 24 |
| AJA305 | B.2.19b | B | Sweden | 55.97744/12.69058 | A.Godhe | 2017-04-07 | ASW | 12-15 | 16 |
| AJA332 | D.1.27a | D | Sweden | 58.33200/16.70583 | A.Godhe, B.Andersson | 2018-05-14 | WC + salt | 8-9 | 8 |
| AJA328 | F.1.2a | F | Sweden | 63.65317/18.95200 (*) | A.Godhe | 2018-03-15 | WC + salt | ~8 | 8 |
| AJA311 | I.3.11a | I | Finland | 60.18000/25.50700 | A.Kremp | 2017-03-09 | WC + salt | 5-6 | 5 |
| AJA313 | J.3.42b | J | Finland | 60.38964/27.37518 (*) | A.Kremp | 2018-04-22 | WC + salt | 4-5 | 8 |
| AJA318 | K.3.3a | K | Estonia | 57.81670/22.28330 | S.Sildever | 2018-05-23 | WC + salt | ~8 | 8 |
| AJA333 | P.2.6a | P | Poland | 54.44778/18.57611 | A.Witkowski | 2018-05-16 | WC + salt | 5-7 | 5 |

In our experiment, we grew eight different *S. marinoi* strains in triplicate for approximately two weeks at three salinities (8, 16 and 24)—a design that included both biological (eight genotypes) and technical replicates (three replicates per strain) (Fig. 1B). Further details about the experimental design, RNA sequencing, and read processing are outlined in the Supplementary Methods. RNA-seq data are available from the Sequence Read Archive (NCBI) under project number PRJNA772794.

We mapped RNA-seq reads against the reference genome of *S. marinoi* strain RO5AC v.1.1 (available from doi 10.5281/zenodo.5266588) with STAR [25], followed by gene-level read quantification with HTSeq [26]. We performed statistical analyses in the R computing environment, using edgeR [27] and stageR [28]. Gene ontology (GO) enrichment was done with TopGO (Over-representation Analysis, ORA) [29] and CAMERA (Gene Set Enrichment Analysis, GSEA) [30]. We obtained functional annotations for all genes using InterProScan [31], KofamKOALA [32], and BLAST+ [33] searches against the Swissprot and Uniprot databases. We detected orthologs of *S. marinoi* genes in other diatom genomes with OrthoFinder [34] and made protein targeting predictions with MitoProt [35], HECTAR [36], SignalP [37], ASAFind [38], and TargetP [39]. Full details on the bioinformatic pipeline, statistical models, and GO enrichment are outlined in the Supplementary Methods, and the gene-level count data and R code used for the statistical analyses are available from Zenodo (doi 10.5281/zenodo.5266588).

### Hypothesis testing

We separately tested two sets of null hypotheses (Fig. 1C). The first set tested whether gene expression was different across the salinity gradient for each genotype separately (Fig. 1C: individual genotypes) and for all genotypes together (Fig. 1C: average of all genotypes), using a total of 27 contrasts (Fig. 1C). Compared to solely testing the average salinity effect, simultaneously accounting for the individual genotypes increases the power to find differentially expressed (DE) genes, as the genotype effect incorporates variability that would otherwise be unaccounted for. The second set of hypotheses tested for an interaction effect between genotype and salinity, i.e., whether there are genotype-specific responses to changes in the salinity (Fig. 1C). Here we defined 84 contrasts, testing each pairwise combination of genotypes within all three salinity combinations (Fig. 1C). We tested the two sets of hypotheses separately using a stage-wise testing procedure in stageR. This allowed us to control the gene-level false discovery rate (FDR) at 5 % when testing multiple hypotheses simultaneously [28].

## RESULTS & DISCUSSION

### General response of Baltic *S. marinoi* to low salinity environments

The inclusion of multiple genotypes in our study design allowed us to characterize both the responses of each individual genotype, and the average response across all genotypes (Fig. 1C). To obtain a general overview of the response of *S. marinoi* to low salinity, we present the results of all three contrasts (8-24, 8-16 and 16-24) together. Unless otherwise indicated, ‘low salinity’ refers to a lower salinity (8 and/or 16) relative to a higher one (16 and/or 24), but does not distinguish between contrasts unless necessary. This was done to gain a general overview of the response to low salinities across the Baltic Sea salinity cline, and because GO enrichment suggested that many of the same processes are enriched in the three salinity contrasts.

Growth reaction norms showed no substantial differences among genotypes and salinity treatments, indicating that all genotypes grew well across a wide salinity range (Fig. 2). RNA-seq reads of all genotypes mapped equally well to the reference genome. In the combined average- and genotype-specific response (Fig. 1C), 7,905 of the 22,440 predicted genes in the *S. marinoi* genome were DE using a 5 % FDR. The number of DE genes in the average response was higher than in any contrast of the individual genotypes (Fig. 3A), with a total of 5,343 DE genes in at least one of three contrasts tested in the average response. This is a result of combining all information from the different genotypes, which resulted in 8×3 replicates for each salinity condition. As a consequence, the average response allowed us to detect many more DE genes, including those with small effect-sizes, and shows the benefit of including more than the standard three replicates used in many transcriptome studies.

**Fig. 2.**
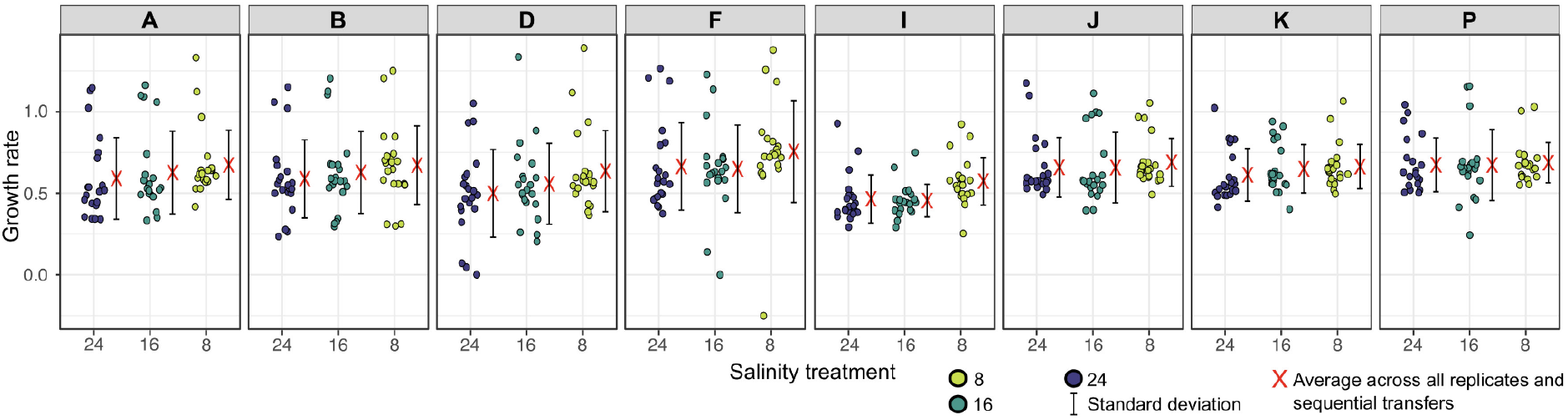
Growth response of Baltic *S. marinoi* in low salinities. Growth rates of the eight *S. marinoi* genotypes examined in this study at three different salinities. The letters in the individual panels correspond with the sampling locations in Table 1 (‘Field ID’) and Fig. 1A. Each point represents a single estimate of the slope of the logarithm of in vivo relative fluorescence against time for each sequential transfer, using a horizontal jitter of points to avoid overplotting.

**Fig. 3.**
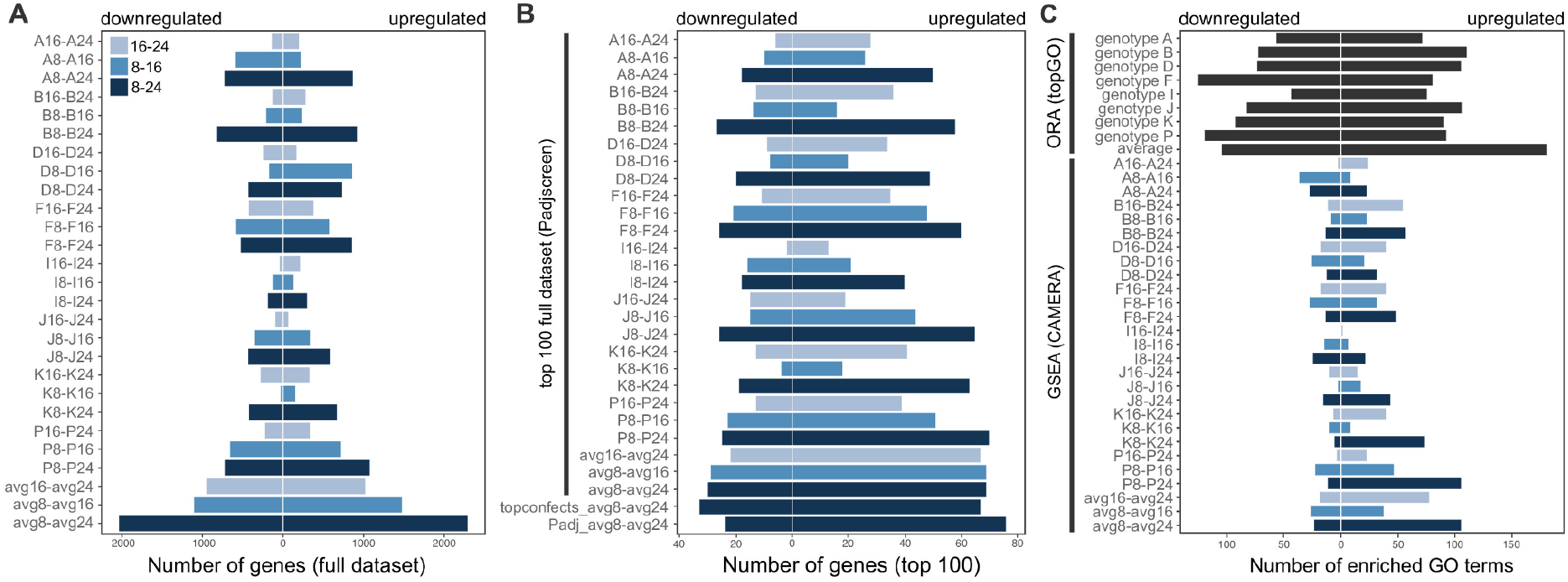
Transcriptome response of Baltic *S. marinoi* to low salinities. **a** Number of DE genes at a 5 % FDR-level in the average and genotype-specific effects. The number of DE genes is indicated separately for each contrast, distinguishing between genes that are up- or downregulated. **b** Direction of DE in the top 100 genes of the average and genotype-specific effects as selected by P-value or logFC. For each contrast in the average and genotype-specific effects (vertical black bar), the direction of DE is indicated for the top 100 genes selected by stageR’s FDR-adjusted P-value of the global null hypothesis (Padjscreen). Thus, although a gene can have a high P-value on a dataset-wide level, it is not necessarily DE in each individual contrast. In addition, we show the top 100 genes selected by logFC (topconfects) and the contrast-specific 5 % FDR-controlled P-value (Padj) for the 8-24 contrast of the average effects, as this contrast showed the greatest number of DE genes in **a**. **c** Number of enriched GO terms. Two types of GO enrichment are shown: Over-representation Analysis, ORA (in topGO) and Gene Set Enrichment Analysis, GSEA (in CAMERA). For ORA, we defined two sets of genes in each genotype/average response for which GO enrichment was performed separately: genes upregulated in low salinities, and genes downregulated in low salinities. Further details on how this selection was performed can be found in the Supplementary Methods. For GSEA, we performed GO enrichment on each contrast of the genotype-specific and average effects separately. Here, the number of up- and downregulated GO terms represents the output classification by CAMERA. The number of enriched GO terms includes Biological Process, Molecular Function and Cellular Component GO terms, prior to removal of redundant GO terms by REVIGO [82] (see Supplementary Methods).

The 8-24 contrasts consistently showed the most DE genes, and the least generally were found in the 16-24 contrast (Fig. 3A). Thus, the largest drop in salinity (24 to 8), and the shift to lower salinity (16 to 8), elicited the greatest transcriptomic responses. When including all DE genes, the number of up- and downregulated genes was comparable within contrasts (Fig. 3A, Suppl. Fig. 1). However, when only taking into account the top 100 genes based on P-value or logFC, substantially more genes were upregulated in low salinities (Fig. 3B), indicating genes with the largest effect sizes or strongest evidence for DE were more likely to be upregulated in low salinities. A similar pattern was seen in the GO enrichment, where a CAMERA analysis found substantially more enriched GO terms that were upregulated in low salinities (Fig. 3C). From these results it is clear that low salinities induce an increased transcriptional response relative to high salinities. In the next paragraphs, we report on several specific pathways that are up- or downregulated in low salinities. Unless otherwise noted, the data in these sections represent the 8-24 contrast of the average response (Fig. 1C).

#### Low salinities trigger profound changes in diatom metabolism

*Skeletonema marinoi* exhibited profound changes in energy metabolism following exposure to low salinity (Suppl. Figs 3-9). In lower salinities, *S. marinoi* experiences (i) significant upregulation of genes from the photosynthetic electron transport chain, Calvin cycle, and chlorophyll biosynthesis (Fig. 4, Suppl. Figs 2, 3A-B, 4A), and (ii) significant downregulation of genes involved in protein ubiquitination, proteolysis, and aerobic respiration (Fig. 4, Suppl. Figs 2, 5A-B). The latter is evidenced by downregulation of the mitochondrial electron transport chain and the TCA cycle, including transcription factor *bZIP14*, which regulates the TCA cycle in diatoms [40] (Suppl. Fig. 5A-B). Thus, the transcriptome suggests relatively more energy is acquired through light reactions, and there is less intracellular recycling of proteins. This is in contrast to another euryhaline diatom, *Thalassiosira weissflogii*, where carbon fixation was not impacted when cells were grown at different salinities [41].

**Fig 4.**
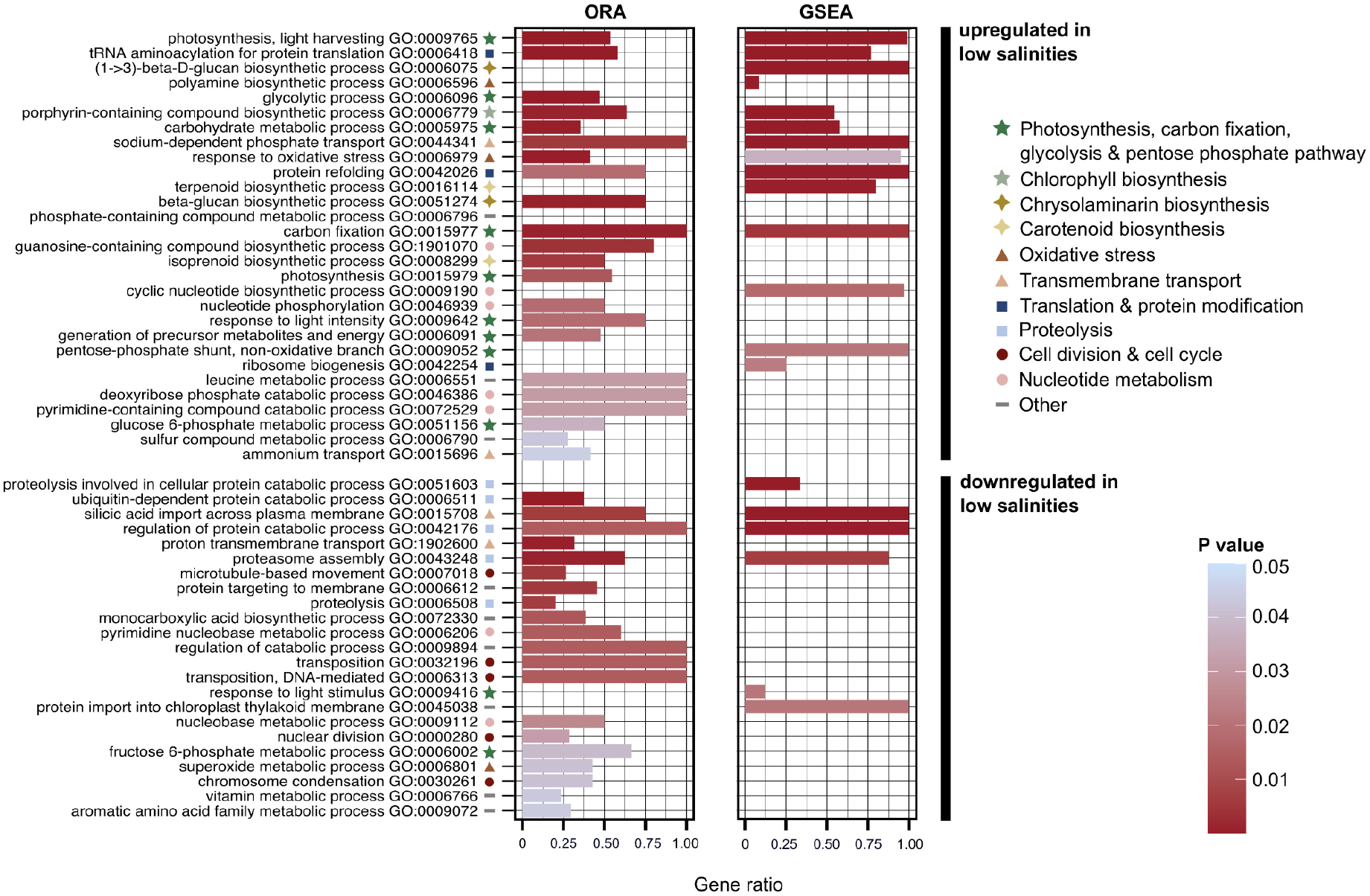
GO enrichment on the average response of *S. marinoi* to low salinities: Biological Process. The results of two types of GO enrichment analyses are shown: ORA (in topGO, Fisher’s exact test, *elim* algorithm) and GSEA (in CAMERA), after removal of redundant terms by REVIGO. For ORA, we classified the total set of DE genes in the average response into two categories, distinguishing between genes that are up- or downregulated in low salinities, regardless of salinity contrast (see Supplementary Methods for more details). For CAMERA, we performed GSEA analyses on each individual contrast separately, showing only the 8-24 contrast in this figure. Barplot height indicates the proportion of genes that are DE with a given GO-term to the total number of genes with this GO-term in the genome of *S. marinoi*. The barplots are colored according to P-value. Within the set of up- and downregulated genes, the GO-terms are ranked from lowest to highest P-value, using the lowest of two P-values from ORA or GSEA. Symbols indicate major categories of cellular processes to which a GO-term belongs. Only Biological Process GO-terms are shown.

Diatoms are known to accumulate storage compounds in unfavorable growth conditions [42, 43], and to modify the fatty acid and lipid composition of their membranes, which alters membrane permeability and fluidity under salinity stress [44]. Here, we observed increased biosynthesis of storage compounds, suggesting that low salinities are stressful to *S. marinoi*, despite their natural occurrence in low-salinity regions of the Baltic Sea. Specifically, biosynthesis of two major storage compounds was upregulated: the polysaccharide chrysolaminarin, i.e. β-1,3- and β-1,6-glucan polysaccharides [45], and fatty acids that might be stored as triacylglycerols [46] (Fig. 4, Suppl. Figs 2, 6A). Four genes involved herein were DE in all genotypes, underscoring that storage compound biosynthesis was a common response in all investigated genotypes. In addition, further evidence from changes in cell membranes came from upregulation of sulfolipids and a glycosyltransferase, which are involved in thylakoid membranes and maintenance of membrane stability, respectively.

We found that tRNA-aminoacylation, translational elongation factors, and many ribosomal proteins were upregulated, which together point to an increase in protein biosynthesis in low salinities (Fig. 4, Suppl. Figs 2, 7). In parallel, protein refolding activity was upregulated (Fig. 4), suggesting proteins need additional stress protection in low salinities. Several transcription factors were also upregulated, including a putative heat stress transcription factor involved in DNA-binding of heat shock promoter elements. Both nuclear division and microtubule-based movements were downregulated (Fig. 4, Suppl. Fig. 2), suggesting less mitosis in low salinities. Although our growth data measured from relative chlorophyll *a* fluorescence do not support a decrease in cell division in low salinities (Fig. 2), upregulation of chlorophyll biosynthesis in these conditions (Suppl. Fig. 4A) suggests that a decrease in mitosis could have been masked by an increase in per-cell chlorophyll-content—an observation that might have important consequences for similar experiments that use chlorophyll *a* as a proxy for growth. In addition, two genes coding for an extracellular subtilisin-like serine protease were upregulated in low salinities, as has also been observed in diatoms in response to other stressors such as copper deficiency [47]. Finally, although activation of transposable elements has been linked to the diatom stress response [48, 49], including *S. marinoi* [50], the majority of genes involved in transposon activity (i.e., transposase, retrovirus-related Pol poly-protein) in the *S. marinoi* genome were not DE, and if DE they tended to be downregulated in low salinities (Fig. 4, Suppl. Fig. 2).

#### Increased nutrient demand in low salinities

Our data suggest that low salinities induce profound changes in nutrient demand, especially of nitrogen, and potentially also highlight preferred nitrogen sources in *S. marinoi*. In fact, many transmembrane transporters for essential nutrients such as nitrogen, phosphorus, molybdate, and sulphate were upregulated in low salinities (Suppl. Figs 5D, 9). Specifically for nitrogen, *Nrt* nitrite/nitrate transporters were highly upregulated, and to a lesser extent transporters for urea and ammonia (Suppl. Figs 5D, 9). Most of the imported nitrogen is directed to the chloroplast, where nitrogen assimilation through ferredoxin-nitrite reductase (*Fd-Nir*) and GSII-GOGAT(Fd) [51] was upregulated (Suppl. Fig. 5D). In parallel, the anabolic part of the urea cycle was upregulated, including carbamoyl phosphate synthase (*unCPS*) (Suppl. Figs 5C-D), suggestive of increased recycling of ammonia and biosynthesis of arginino-succinate or arginine. Higher nutrient demands in low salinities likely reflect increased biosynthesis of nitrogen-rich compounds such as amino acids and polyamines, essential for protein biosynthesis and the stress response, respectively [52, 53]. In parallel, we detected a high number of DE transporters for amino acids and polyamines, most of which were upregulated in low salinities (Suppl. Fig. 9). Taken together, it appears that *S. marinoi* requires more nutrients in low-salinity environments to support increased demand for proteins and other compounds. Consequently, *S. marinoi* might struggle to maintain growth in low salinities under nutrient-deplete conditions, suggesting that increasing nutrient loads in low-salinity regions of the Baltic Sea might benefit *S. marinoi* [54, 55]. In contrast, silicic acid transporters were strongly downregulated in low salinities, and this downregulation was most evident in the 16-24 salinity contrast (Suppl. Fig. 9). This could reflect decreased cell division and/or changes in cell wall silicification.

#### A diverse response to oxidative stress in low salinities

Several lines of evidence suggest that *S. marinoi* experiences increased oxidative stress at low salinities, as multiple mechanisms to deal with reactive oxygen species (ROS) were upregulated in low salinities. This included genes involved in glutathione metabolism, ascorbate peroxidases, catalases, and peroxiredoxin (Suppl. Fig. 6B). The xanthophyll cycle, which plays a critical role in protection from oxidative stress due to excess light and ROS generated by other stressors [56], was also distinctly upregulated (Suppl. Fig. 4B). Carotenoids for the xanthophyll cycle were produced primarily through the non-mevalonate pathway, which was also upregulated in low salinities (Suppl. Fig. 4B).

We found upregulation of polyamine biosynthesis from ornithine via ornithine decarboxylase (Suppl. Figs 5C-D). Polyamines play a complex role in abiotic stress responses, including salinity stress in land plants, by increasing the activity of antioxidant enzymes, triggering the stress signal transduction chain and serving an osmolyte function [53]. In diatoms, polyamines increase in response to both heat and salinity stress [57, 58], and our data suggest a similar role in salinity acclimation.

Violaxanthin-de-epoxidase (xanthophyll cycle), and two genes involved in polyamine biosynthesis were DE in all genotypes (Fig. 5), underscoring the highly conserved nature of the oxidative stress response in *S. marinoi*. In addition, vitamin B6-binding activity was upregulated in low salinities. Vitamin B6 is a vital cofactor of various enzymes, and plays a role in ROS deactivation in land plants [59]. However, the response of *S. marinoi* to oxidative stress is complex, as several other genes involved in ROS elimination were downregulated in low salinities (Suppl. Fig. 6B), including superoxide dismutase (SOD), which is a first line of defense against ROS in land plants and macroalgae [60, 61].

**Fig. 5.**
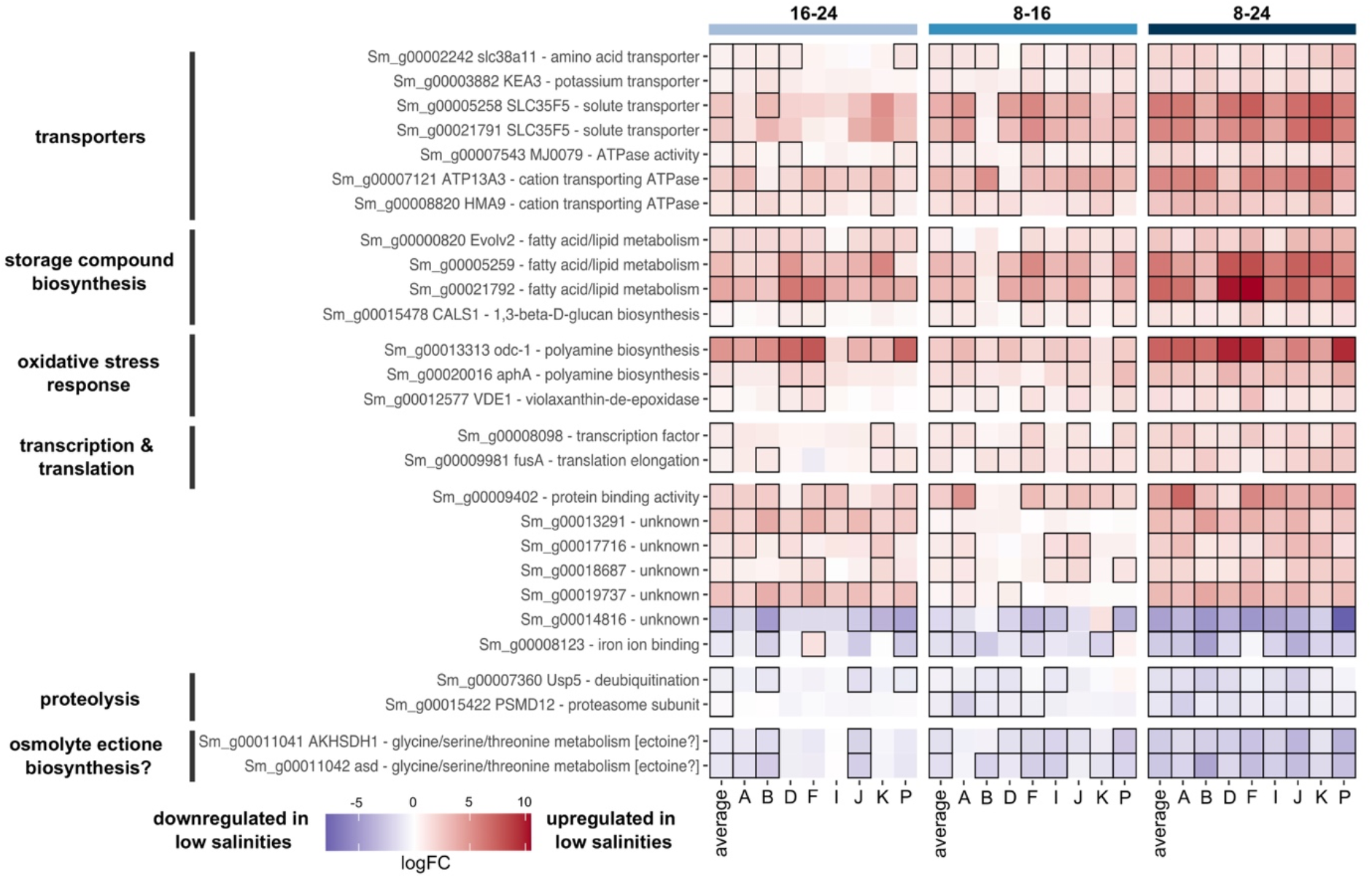
Set of genes that are DE in at least one contrast of each genotype: the core response. The heatmap shows logFC values for the individual genotypes and average response of the 27 core response genes. The three salinity combinations are indicated on top of the figure. Contrasts that were significant are outlined in black. Rownames specify gene names and functional annotation. When DE, all genes are consistently up- or downregulated in low salinities in each genotype, with the exception of gene *Sm_g00008123*.

#### Osmotic stress response

*Skeletonema marinoi* produces a variety of osmolytes. These are small organic molecules that help mitigate hyperosmotic stress typical of marine environments [62–65]. Consequently, a decrease in salinity should trigger a drop in osmolyte biosynthesis. Indeed, in low salinities, *S. marinoi* adjusted intracellular osmolyte concentrations to hypoosmotic conditions by downregulating biosynthesis of the osmolyte DMSP and upregulating breakdown of the osmolyte taurine via taurine dioxygenase (Suppl. Fig. 6B). Although most genes involved in the diatom DMSP pathway remain unknown, *S. marinoi* has a homolog of *TpMMT*, a methyltransferase that catalyzes a key reaction in DMSP biosynthesis in diatoms [64]. This gene was strongly downregulated in low salinities (Suppl. Fig. 6B). Putative *BADH* and *CDH* genes involved in the biosynthesis of the osmolyte betaine from choline [62] were present but not DE. Betaine can also be synthesized from glycine by *TpGSDMT [62]*, and a possible homolog of this gene was downregulated in all genotypes (Suppl. Fig. 6B). Genes involved in proline metabolism were generally slightly upregulated (Suppl. Fig. 5D), suggesting proline is the preferred osmolyte in low salinities.

Two genes that were strongly downregulated in low salinities in all genotypes could be involved in the biosynthesis of a fifth osmolyte: ectoine (Fig. 5, Suppl. Fig. 8). These genes encode a bifunctional aspartokinase/homoserine dehydrogenase (*Sm_g00011041*) and an aspartate-semialdehyde dehydrogenase (*Sm_g00011042*) (Fig. 5, Suppl. Fig. 8). Both are involved in the early steps of the glycine/threonine/serine pathway, and convert aspartate into aspartate-semialdehyde and/or homoserine. The genome of *S. marinoi* contains several other homologs of both genes. When DE, these homologs show opposite expression patterns to the aforementioned genes: they are upregulated in low salinities, following the same expression patterns as other genes in this pathway (Suppl. Fig. 8). Peptide targeting predictions showed that reactions in this pathway are compartmentalized across the chloroplast, cytoplasm, and mitochondria, presumably allowing *S. marinoi* to run opposite reactions simultaneously while avoiding futile cycles (Suppl. Fig. 8). Given their expression patterns and compartmentalization, *Sm_g00011041* and *Sm_g00011042* are likely not involved in conventional amino acid biosynthesis. Instead, one of their products, aspartate-semialdehyde, is a known precursor for the osmolyte ectoine in bacteria (Suppl. Fig. 8) [66], and elevated levels of aspartate-semialdehyde dehydrogenase have been detected in bacteria occupying high salinities [67]. Recently, marine diatoms were found to both biosynthesize and import ectoine of bacterial origin [68]. Several *S. marinoi* genes may be homologous to the bacterial ectoine genes *ectA*, *ectB*, and *ectC* that convert aspartate-semialdehyde to ectoine. However, low sequence similarity (max. 47.8 %), and lack of downregulation in low salinities, raises doubt about whether those genes are responsible for ectoine biosynthesis in *S. marinoi*. Furthermore, all putative homologs received annotations different from ectoine-related genes in Swissprot/Uniprot, suggesting they serve different functions from aforementioned ectoine genes. It is possible that diatoms have other unknown genes involved in ectoine biosynthesis, or alternatively, diatoms might provide ectoine precursors (e.g., aspartate-semialdehyde) to extracellular bacteria that synthesize and return ectoine to the diatom. Such exchange of metabolites has previously been shown to occur in diatom–bacteria interactions [69]. Our expression data are consistent with both scenarios (Suppl. Fig. 8), and suggest ectoine might be an important osmolyte in diatoms.

Responses to osmotic stress also include shifts in cation import and export, such as sodium and potassium [70]. Here, most identified potassium and sodium channels were either slightly upregulated or not DE (Suppl. Fig. 9), and two detected aquaporins were either up- or downregulated in low salinities. Six transporters for potassium, amino acids or unknown cations/solutes were DE in all genotypes, often with large effect-sizes (Fig. 5), highlighting the conserved nature of this response in *S. marinoi*.

### Genotype-specific data reveal intraspecific variation and a conserved core response to low salinity

The genotype effect in our dataset overshadowed the salinity effect, indicating that genotypes differed substantially in their response to low salinities. This was evidenced by a multidimensional scaling plot and poisson-distance heatmap in which samples clustered primarily by genotype rather than salinity (Fig. 6). Specifically, 1,791 genes were uniquely DE in only one genotype (Suppl. Fig. 10). The number of uniquely DE genes ranged from 103 (genotype I) to 317 genes (genotype A) (Suppl. Fig. 10).

**Fig. 6.**
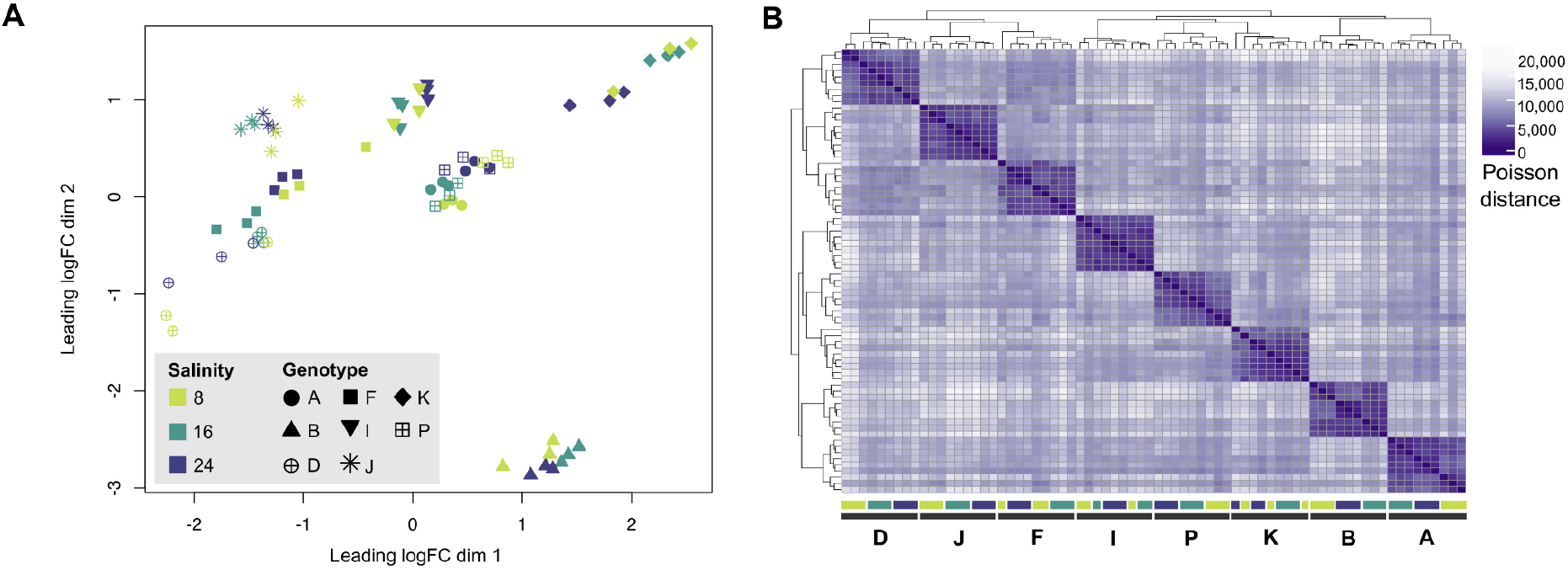
Intraspecific variation in the response of Baltic *S. marinoi* to low salinities. **a** Multidimensional scaling (MDS) plot for the full dataset, showing that samples cluster primarily by genotype rather than salinity. Distances between the samples are based on logFC changes in the top 500 genes. **b** Poisson-distance heatmap of the full dataset. Colored bars below the heatmap indicate the position of samples belonging to different genotypes and salinities (Fig. 1 a), showing that samples of different genotypes cluster together.

We defined a core response to low salinities by selecting genes that are DE in at least one contrast of each genotype, which resulted in a set of 27 shared genes that are DE in each genotype (Figs 1C, 5). Obtaining this set of shared genes required subsetting the full set of DE genes (Fig. 1C), so the 5 % FDR for these 27 genes could not be guaranteed. However, the core response genes were characterized by a combination of high logFC and low P-values (Suppl. Fig. 1), thus providing strong evidence for DE in each genotype. These genes are also among the top DE genes in the full dataset (Fig. 1C: blue section): 13 overlapped with the top 25 DE genes as ranked by stageR’s FDR-adjusted P-value of the global null hypothesis (Padjscreen), 22 were part of the top 100 DE genes, and all were detected within the top 225 DE genes. Core response genes upregulated in low salinities were involved in key processes previously identified in the average response, including transport of amino acids and cations, storage compound and polyamine biosynthesis, ROS deactivation, and transcription and translation (Fig. 5). By contrast, core response genes that were downregulated in low salinities were involved in proteolysis or the putative biosynthesis of the osmolyte ectoine (Fig. 5). Increasing the number of technical replicates would likely enlarge the set of core response genes, as higher replicate numbers are bound to improve detection of DE genes, especially those with small effect sizes [71]. Our set of ‘core response’ genes is thus not exhaustive, but gives a first indication of which genes are likely to be part of a conserved ancestral response to low salinity environments in *S. marinoi*.

### Interaction-effects reveal differences among genotypes in their response to low salinity

A total of 3,857 genes showed interaction effects (Fig. 1C), i.e., significantly different expression patterns between genotypes with a 5 % FDR. Of these, 2,820 differed between genotypes in the magnitude of their response to low salinities, whereas far fewer (1,037) differed in the direction of their response (Fig. 1C). However, 92 of the top 100 genes with interaction effects (ranked by stageR’s FDR-adjusted P-value of the global null hypothesis, Padjscreen) differed in the direction of their response (Suppl. Fig. 11). Thus, although more genes overall showed differences in the magnitude of DE, those with differences in direction of DE dominated the top set of interaction-effect genes.

The two classes of interaction-effect genes were enriched for different processes (Fig. 7, Suppl. Fig. 12). Genes that differed significantly across genotypes in the magnitude of their response were enriched for many of the same key processes identified in the average response, including photosynthesis, glycolysis, and the biosynthesis of chlorophyll, carotenoids, and fatty acids (Fig. 7, Suppl. Fig. 12). This suggests that there is considerable variation among *S. marinoi* genotypes in the strength of the salinity response. By contrast, genes that differ significantly between genotypes in the direction of their response were enriched for processes involving transcription regulation, peroxidase activity, aerobic respiration, and nitrogen metabolism, including urea transmembrane transport (Fig. 7), suggesting that genotypes differ, at least partially, in the invoked strategies to cope with low salinity. The top 100 set of interaction-effect genes further suggested genotypes differ in the direction of their response for cell wall and calcium-binding messenger proteins, as well as heat shock proteins/chaperones (Suppl. Fig. 11), highlighting intraspecific differences in how salinity stress affects protein function and cell-signalling pathways. For example, Ca^2+^-signalling is involved in osmotic sensing in diatoms [72], suggesting *S. marinoi* genotypes differ in how they perceive and respond to osmotic stress.

**Fig. 7.**
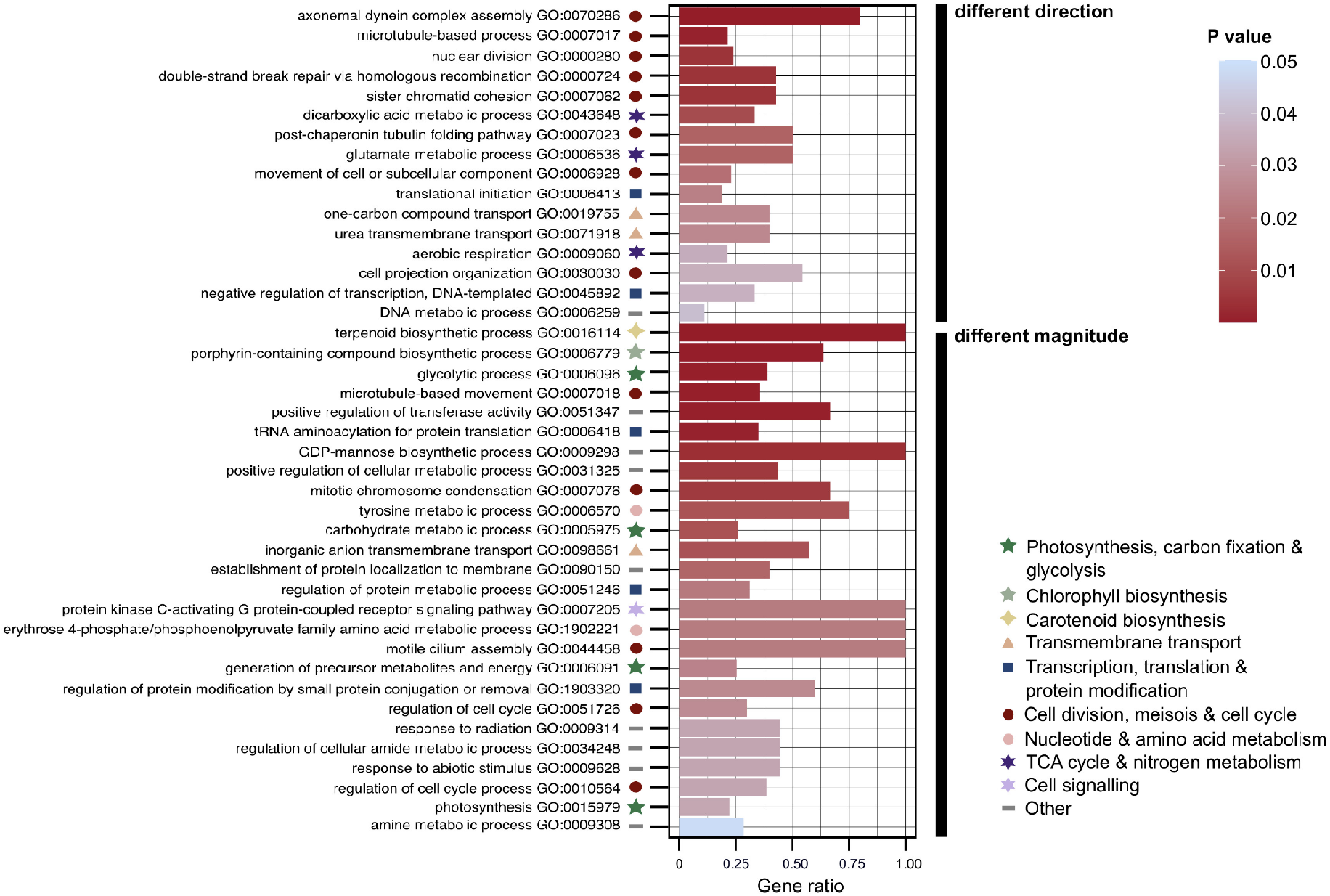
GO enrichment of the interaction-effects: Biological Process. The barplot visualizes the significant GO terms retrieved by ORA (topGO, Fisher’s exact test, *elim* algorithm) after removal of redundant GO terms by REVIGO. Two sets of GO enrichment were carried out which distinguished between genes that differ significantly between genotypes in the direction or magnitude of their response to low salinities. Barplot height indicates the proportion of genes that are DE with a given GO-term to the total number of genes with this GO-term in the genome of *S. marinoi*. The barplots are colored, and the GO terms ranked, according to P-value. Symbols indicate major categories of cellular processes to which a GO-term belongs. Only Biological Process GO-terms are shown.

Both classes of interaction-effect genes contained GO terms related to translational activity, cell cycle progression, mitosis, and meiosis. In fact, two interaction-effect genes that show differences in the direction of their response across genotypes coded for meiotic recombination protein *SPO11-2* (Suppl. Fig. 11), which is thought to be expressed exclusively during meiosis [73, 74]. Sexual reproduction in *S. marinoi* can be induced by shifts from low to higher salinity, e.g., 6 to 16 [73], but our data suggest genotypes differ in the optimal salinity shift to induce meiosis. However, meiosis likely never dominated during our experiment as growth rates were relatively stable over time (Fig. 2), and gametes or auxospores were not observed. Although these observations could reflect intraspecific differences in growth and sexual reproduction related to salinity, we cannot rule out that genotypes were at different stages of their cell cycle when RNA was harvested.

The observed intraspecific variation in gene expression likely reflects an important mechanism which allows *S. marinoi* to grow throughout the Baltic Sea. Diatom populations can harbour high levels of both genotypic and phenotypic variation [23, 75–78]. Our finding of high levels of variation in gene expression is a natural extension of this observation. Our study design did not allow testing whether the observed intraspecific variation is related to the natural salinities at which the different genotypes occur, as this would require sampling of multiple genotypes within populations. Nevertheless, visual comparison of gene expression patterns did not show consistent differences across low- (D, F, I, J, K, and P) and high-salinity (A and B) populations (e.g., Suppl. Figs 3-9), nor did those populations cluster separately in the MDS-plot (Fig. 6A). This suggests that if signals of local adaptation along the Baltic salinity cline [22] are due in part to differences in gene expression between high- and low-salinity populations, then those differences are subtle. In any case, substantial intraspecific variation in gene expression patterns in *S. marinoi* exists and is likely to be a crucial factor for survival, acclimation, and adaptation to a dynamic environment such as the Baltic Sea, where in addition to a salinity gradient, marked gradients and seasonal fluctuations in nutrients and temperature also occur [55]. The strong degree of variation in gene expression within this population increases the chance that at least some cells can survive rapidly fluctuating, potentially adverse, conditions in the short term. Assuming some of this variation is heritable, variable gene expression can also enable long-term evolutionary adaptation by providing targets for natural selection [8, 79]. In that sense, it may be that the observed variation in gene expression is maintained by balancing selection.

## CONCLUSION

Our study design, in which transcriptome data from eight genotypes were combined into one analysis, allowed for a holistic view of the response of *S. marinoi* to low salinity conditions in the Baltic Sea, the world’s largest brackish water body. Transcriptome studies often include technical replicates of a single genotype, but an increasing number of studies [49, 80] show that experiments without biological replicates are unlikely to be generalizable, as different genotypes can exhibit markedly different patterns of gene expression. Here, inclusion of both technical and biological replicates allowed us to pinpoint both conserved and variable responses to low salinity.

Despite long-term presence of *S. marinoi* in brackish habitats in the Baltic Sea, our transcriptome data suggest that when Baltic *S. marinoi* is exposed to low salinities that mimic the natural Baltic salinity gradient, the diatom experiences elevated stress and increased energy and nutrient demands to maintain homeostasis. This suggests that *S. marinoi* has not fully adapted to low salinity regions of the Baltic Sea since it first colonized the Baltic some 9 000 years ago. Our analyses further revealed substantial levels of intraspecific variability in the response of *S. marinoi* to low salinities. Although our experimental design did not allow us to link this variability to the natural salinity of the source population, it highlights an important source of biological variation in natural populations of *S. marinoi* and presumably other diatoms as well. Variable gene expression could be an important mechanism at which diatoms respond and adapt to environmental change, and ultimately diversified in a wide range of marine, freshwater, and terrestrial habitats worldwide [4, 81].

## ACKNOWLEDGEMENTS

This work was supported by grants from the Simons Foundation (725407, EP and 403249, AJA) and National Science Foundation (1651087, AJA). EP also benefited from postdoctoral fellowships from Fulbright Belgium and the Belgian American Education Foundation (B.A.E.F.). K.V.d.B. is a FWO junior postdoc fellow (project 1246220N) and previously benefited from a B.A.E.F. grant. We are grateful to Sirje Sildever, Björn Andersson and Andrzej Witkowski for sample collections. We thank Wade Roberts, Quinten Bafort, and Gust Bilcke for providing advice on the data analysis.

## COMPETING INTERESTS

The authors declare they have no conflict of interest.

## Notes

### Competing Interest Statement

The authors have declared no competing interest.

