## Supplementary Information for "Genotype-specific transcriptional responses overshadow salinity effects in a marine diatom sampled along the Baltic Sea salinity cline"

This Supplementary Information file includes the following items:

- Supplementary Methods
- Supplementary Figures 1 - 13
- Supplementary References

### SUPPLEMENTARY METHODS

#### Genotyping *S. marinoi* strains

We genotyped all strains obtained from the sediments for D1-D2 LSU rDNA (28S) to ensure they belonged to the species *S. marinoi*. To this end, we harvested cells by centrifugation and flash freezing in liquid nitrogen after which they were stored at -80 °C until DNA extraction. Frozen cells were broken with 1.0 mm glass beads (Biospec Products, OK, USA) by shaking the tubes for 30 seconds at 3 000 OSM in a Minibeadbeater (Biospec Products, OK, USA). We extracted DNA with the DNeasy® Plant Mini Kit (Qiagen, Hilden, Germany), following the manufacturer's instructions. D1-D2 28S (~600 nt) was amplified by PCR using primers D1R and D2C [1]. PCR reactions consisted of 1.0-5.0 µL DNA extract, 6.5 µL of Failsafe Buffer E (Epicentre Technologies, WI, USA), 0.5 µL of each primer (20 µM stocks), and 0.5 units Taq polymerase, using adjustment with $ddH_2O$ to a final volume of 25 µL. We carried out PCR reactions in a T100 Thermal Cycler (Bio-Rad, Hercules, CA, USA) using the following PCR program for all reactions: initial step at 95 °C for 5 min, followed by 36 cycles (95 °C for 50s, 56 °C for 60s and 72 °C for 60s), and ending with a final elongation step at 72 °C for 5 min. The resulting PCR products were purified by treatment with an Exonuclease I (Exo) and Shrimp Alkaline Phosphatase (SAP) protocol that included addition of 0.25 µL Exo, 1.75 µL $ddH_2O$, and 1.o µL SAP per 25 µL PCR product, followed by heating for 30 min at 37 °C and a 15-min termination step at 80 °C. Only forward strands were sequenced at Eurofins Genomics (Louisville, KY, USA). We edited the sequenced chromatograms with Sequencher v5.1 (Gene Codes Corporation, Ann Arbor, MI, USA). In order for a strain to be included in our study, its D1-D2 28S sequence had to be identical to the 28S sequence of *S. marinoi* strain RO5AC for which a genome-sequence is available.

#### Experimental setup, RNA extraction and sequencing

The salinity experiment was run in May-June 2019, at which point the investigated strains were between 12 and 26 months old (Table 1). Throughout the experiment, we used ASW medium with constant nutrient levels, but varying salt concentrations depending on the treatment. The experiment was carried out on a single shelf of a Percival incubator (Percival Scientific, IA, USA) at 12 °C and a 12:12 light:dark light regime at 30 µmol $m^{-2}$ $s^{-1}$ light intensity. During one month, all 72 cultures were grown simultaneously in 4 mL tubes divided among three racks, and all tubes were pseudorandomized daily to avoid potential batch effects. We measured relative chlorophyll *a* fluorescence of each tube daily using a Trilogy fluorometer (Turner Designs, CA, USA), allowing us to monitor growth. Every four days, we transferred the cultures into fresh media, with the first re-inoculum taking place after seven days. Cultures were reinoculated six times, allowing them to maintain exponential growth throughout the experiment. For each reinoculum, we brought the cell densities in all tubes to the same level, using the relative fluorescence data as guideline. With each reinoculation starting from day 11 ($2^{nd}$ reinoculation), we harvested cells for RNA extraction by centrifugation and freezing at -80 °C. To obtain sufficient RNA for sequencing, total RNA was extracted from cells harvested from two serial reinoculations. We randomized all samples over five batches, and extracted RNA with the Qiagen RNeasy plant mini kit. RNA quality was measured using an Agilent TapeStation 2200 (Agilent, CA, USA). Indexed RNA-seq libraries were prepared with the KAPA mRNA HyperPrep library kit using the standard protocol but with half reaction volumes, after which library quality and quantity was assessed with the TapeStation. All libraries were pooled and sequenced together on a single lane of an Illumina HiSeq4000 (2 x 100 paired-end reads) at the University of Chicago Genomics Facility. An average of 9.6±2.4 million reads per sample was sequenced.

**Read trimming and mapping of RNA-seq data**

We performed quality-control of the raw reads using FastQC v0.11.5 [2], after which we applied Ktrim v1.1.0 [3] for adapter removal and quality-trimming, using default settings (baseline phred score, -p 33; minimum quality score, -q = 20; minimum read size after trimming, -s = 36). We subsequently mapped the reads against the reference genome of *S. marinoi* strain RO5AC v.1.1 (available from doi 10.5281/zenodo.5266588) using STAR v.2.7.3a [4] with default settings, except for intron size. For the latter, we used *-alignIntronMin 4* and *-alignIntronMax 17105*. These values were based on the *S. marinoi* genome assembly used for our analysis. We used the uniquely mapping reads for gene-level read quantification in HTSeq v0.11.3 [5] via *union* mode. The output of HTSeq was imported in R v4.0.2 (R Core Team, 2020) for further statistical analysis.

**Functional annotation of *S. marinoi* genes**

We obtained functional annotations of the complete set of *S. marinoi* genes using various approaches. First, we used BLAST+ v2.6.0 [6] to run sequence similarity blastp searches of all *S. marinoi* proteins annotated in the genome against the Swissprot (download June 2020) and Uniprot databases (download June 2019), and retained the best hit using a maximum e-value limit of 1e-6. Second, we ran InterProScan v5.36-75.0 [7] against the InterPro collection of protein signature databases, including Gene Ontology (GO) resources, Pfam domains, PRINTS, PANTHER, SMART, SignalP_EUK, and TMHMM. Third, we obtained KEGG pathway annotations via the KofamKOALA web server v2020-08-04 (KEGG release 95.0) [8]. At last, we predicted protein targeting for a subset of genes involved in nitrogen metabolism, the pentose phosphate pathway, the Calvin cycle, the TCA cycle, glycine/serine/threonine metabolism, and glycolysis and gluconeogenesis using a method slightly adapted from [9, 10]. More specifically, we used the software programs MitoProt [11], HECTAR v1.3 [12], SignalP-3.0 [13], ASAFind [14] and TargetP-2.0 [15] to predict protein localization to the mitochondria, chloroplasts or cytoplasm. If a protein had predicted plastid-peptides in HECTAR, SignalP and ASAFind it was classified as plastid-targeted, whereas if it was targeted to the mitochondria by any two of MitoProt, HECTAR or TargetP it was classified as mitochondria-targeted. As no endoplasmic reticulum-targeted signal peptides were detected, all proteins with no clear targeting were assumed to be active in the cytoplasm. If conflicting results were obtained, proteins were classified as having dual or uncertain targeting.

**Homology search in the *S. marinoi* genome**

In order to find orthologs of the *S. marinoi* genes in other diatoms for which a genome is available, we ran OrthoFinder v2.2.6 [16] in DIAMOND mode including nine other diatom proteomes, i.e. *Cyclotella cryptica*, *Cyclotella nana*, *Fistulifera solaris*, *Fragilariopsis cylindrus*, *Nitzschia* sp., *Phaeodactylum tricornutum*, *Pseudonitzschia multiseries*, *Pseudonitzschia multistriata*, and *Seminavis robusta* [10, 17–23]. In addition, we used BLAST+ with an e-value limit of 1e-6 to search for homologs of a series of bacterial genes in the *S. marinoi* genome. This was done for the enzymes *ectA*, *ectB* and *ectC* which are involved in the ectoine pathway in bacteria. To this end, all available sequences of bacterial enzymes were downloaded from Uniprot, and for each the best hit was retained in BLAST+.

**Differential expression analysis**

Using the R-package edgeR v3.34.0 [24], we filtered the gene-level counts to only include genes that have at least one count per million (CPM) in at least three samples. TMM normalization (i.e. weighted trimmed mean of the log expression ratios) was used to eliminate technical variation due to library size and composition [25]. We created a multidimensional scaling (MDS) plot based on Euclidean distances on the gene expression profiles of the pairwise top 500 genes using the R-package limma v.3.48.0 [26]. Subsequently, we used edgeR to fit a quasi-negative binomial generalized linear model (GLMs) [27] for every gene using the glmQLFit function with a group-model design that included each genotype-treatment combination. Hypothesis testing was performed using F-tests with the glmQLFTest function. Due to the multiplicity of tests performed for each gene, we used stage-wise testing in stageR v1.14.0 [28]. Specifically, in the screening stage of stageR, we tested the global null hypothesis, i.e. the null hypothesis over all contrasts together, on a 5 % FDR using the Benjamini-Hochberg correction [29]. We then tested all contrasts separately in the confirmation stage, only including genes that were significant in the screening stage, and Holm's method [30] was used to control the within-gene family-wise error rate (FWER) on the adjusted FDR-level of the screening stage, altogether controlling the gene-level FDR at 5 % [28, 31]. Sets of top genes of interest for the average and genotype-specific responses were selected using stageR's FDR-adjusted P-value of the global null hypothesis (Padjscreen). In addition, we selected top gene sets for individual contrasts using contrast-specific FDR-controlled P-values, as well as logFC values via the Topconfects method [32].

**GO enrichment**

For each significant gene in the confirmation stage of the stage-wise testing analysis, we extracted Gene Ontology (GO) terms from the InterProScan results and subdivided these into the three main GO categories: Biological Processes (BP), Molecular Function (MF), and Cellular Component (CC). We performed GO enrichment on the results of the individual genotypes and the average response (first set of hypotheses), and the interaction-effects (second set of hypotheses). For the individual genotypes and the average response, we performed GO enrichment for each of the 27 contrasts in CAMERA [33] as implemented in edgeR, separately for each main GO category. In addition, we defined two main categories of genes in the individual genotype and average responses, separately for each genotype and the average response: (i) upregulated in low salinities, and (ii) downregulated in low salinities. We assigned genes to these two categories based on their expression patterns in the three salinities, visualized using the TMM normalized logarithm of the average expression of each gene in function of salinity, i.e. the log fitted values of the glmQLFit output (Suppl. Fig. 13). Genes that were up- or downregulated in intermediate salinities were discarded as these sets were too small to perform robust GO enrichment analysis (Suppl. Fig. 13). For each of the two categories, GO enrichment was performed using Fisher's exact test and the *elim* algorithm in the R-package TopGO v2.44.0 [34], separately for each main GO category.

For the interaction effects, genes were assigned to two sets: (i) genes responding in similar directions across genotypes, and (ii) genes responding in different directions across genotypes. Set (i) also included genes that were only DE in one genotype. To determine to which category a gene belonged, we used the logFC values of DE genes from the contrasts of the individual genotypes (first hypothesis, Fig. 1C), i.e., when for a given gene all logFC values of DE contrasts of the genotypes were positive or negative, a gene belonged to category (i), whereas if both positive and negative logFC values were detected, a gene was assigned to category (ii). Genes that were not DE in any contrast of the individual genotypes were assigned to (i). Twenty-four genes of (ii) were uniquely DE in one genotype, but showed both positive and negative logFC values in different contrasts, indicating differences in the direction of DE between different contrasts within a genotype, but not across genotypes. These genes were removed from (ii) and added to (i) prior to GO enrichment, because they represent a difference in magnitude of the response across genotypes. This ensured (ii) only included genes that differed in the direction of DE across genotypes, whereas (i) included all other genes with interaction-effects which exhibited differences in magnitude, but not direction, across different genotypes. When assigning genes, we did not distinguish between the three different sets of salinity contrasts (16-vs-8, 24-vs-16 and 24-vs-8) because this would have reduced the set of DE genes too much to allow for robust GO enrichment. Subsequently, we performed GO enrichment of the two sets of interaction-effect genes in TopGO as outlined above. For all TopGO analyses, the selected set of genes was compared to the full set of genes in the genome of *S. marinoi*. For both the TopGO and CAMERA results, we used REVIGO [35] to summarize significantly ($P < 0.05$) enriched GO terms, by means of the SimRel score [36].

**SUPPLEMENTARY FIGURES**

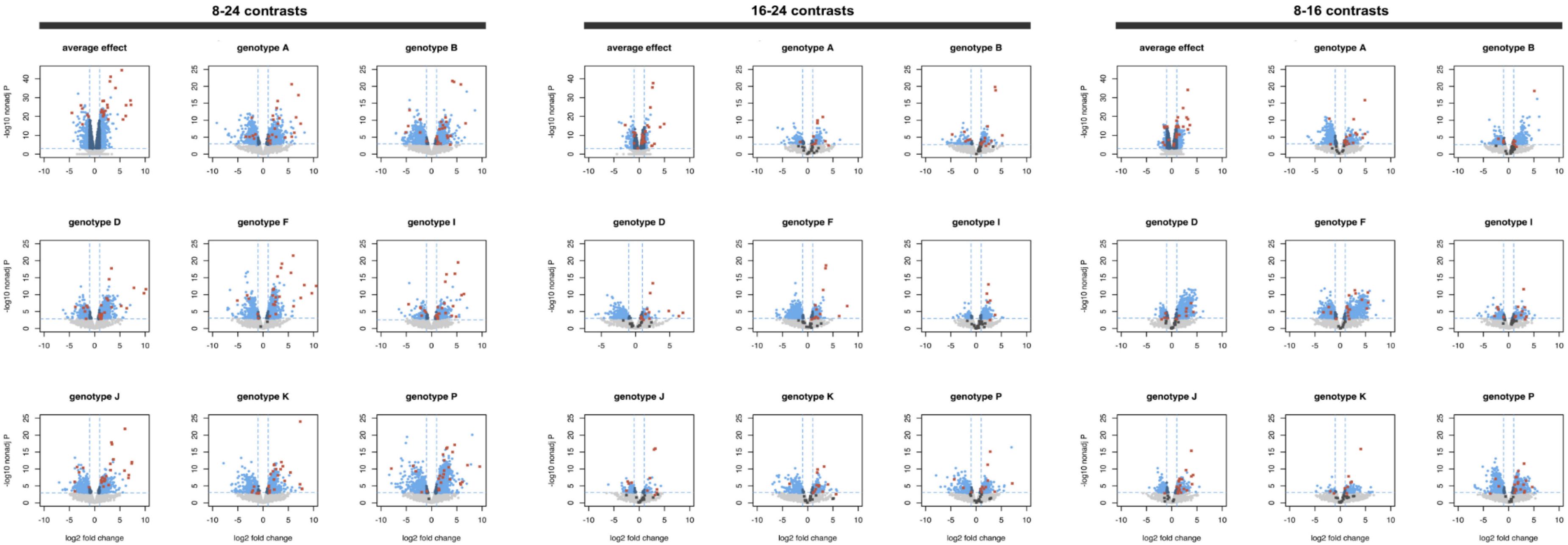


**Suppl. Fig. 1. Volcano plots of all the contrasts for the average and genotype-specific effects.** The plots depict the logFC versus the log10 5 % FDR-adjusted P-values. All values above the horizontal dotted blue lines are significant, and all values below this line are not significant. DE core response genes are indicated with red squares. Dark grey squares indicate core response genes that are not significant in a given contrast.

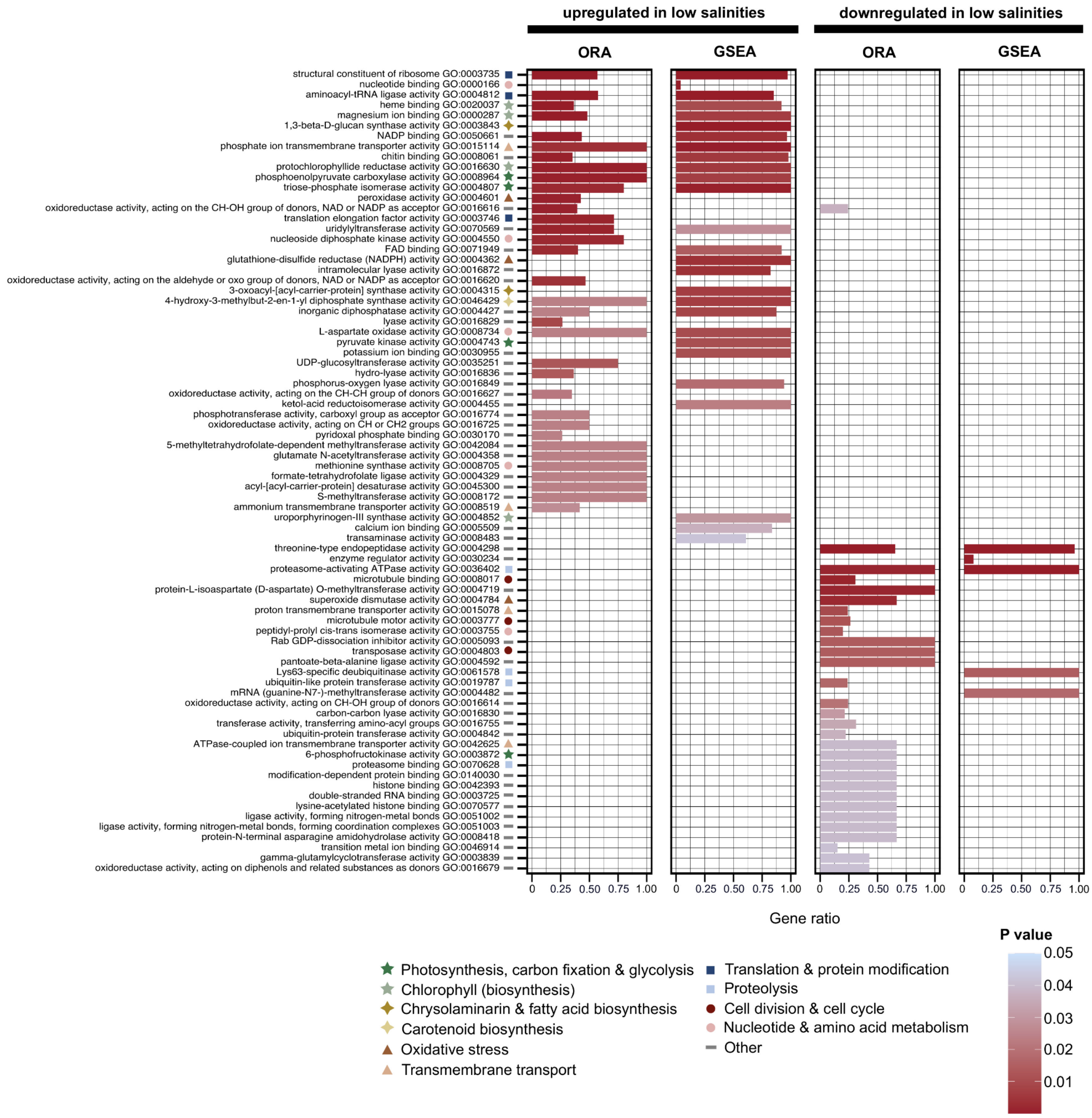


**Suppl. Fig. 2. GO enrichment on the average response of *S. marinoi* to low salinities: Molecular Function**. The results of two types of GO enrichment analyses are shown: ORA (in topGO) and GSEA (in CAMERA), after removal of redundant terms by REVIGO. For ORA, we classified the total set of DE genes in the average response in two categories, distinguishing between genes that are up- or downregulated in low salinities, regardless of salinity contrast (see Supplementary Methods for more details). For CAMERA, we performed GSEA analyses on each individual contrast separately, showing only the 8-24 contrast in this figure. Barplot height indicates the proportion of genes that are DE with a given GO-term to the total number of genes with this GO-term in the genome of *S. marinoi*. The barplots are colored according to P-value. Within the set of up- and downregulated genes, the GO-terms are ranked from lowest to highest P-value, using the lowest of two P-values from ORA or GSEA. Symbols indicate major categories of cellular processes to which a GO-term belongs. Only Molecular Function GO-terms are shown.

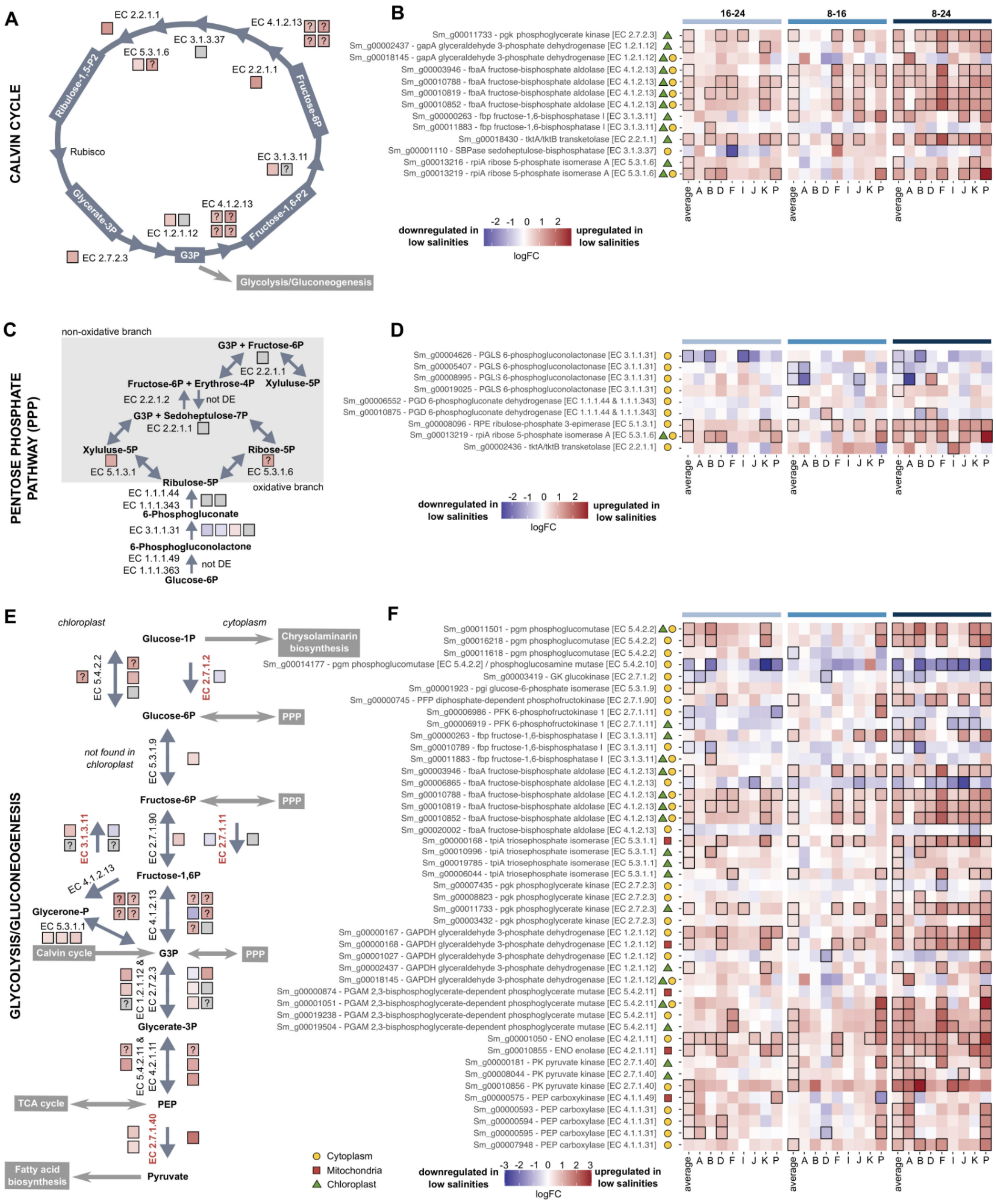

**Suppl. Fig. 3. Gene expression of genes involved in carbohydrate metabolism, showing pathways and heatmaps on gene expression. a, b** Calvin cycle. **c, d** Pentose phosphate pathway. **e, f** Glycolysis/gluconeogenesis. All heatmaps (**b, d, f**) show logFC values of the 16-24, 8-16 and 8-24 contrasts of the average response and the eight genotypes. All visualized genes are DE in at least one contrast. Contrasts that were significant are outlined in black. Symbols next to the gene names in the heatmaps denote protein targeting. In case the protein targeting was unclear, multiple symbols are used. Each colored square in the pathway figures (**a, c, e**) corresponds with a single gene, and is colored according to the logFC values of the 8-24 contrast of the average response. Squares filled with a question mark indicate proteins with unclear targeting. In panel (**e**), the genes left and right of the pathway arrows are targeted to the chloroplast and cytoplasm, respectively. In case protein targeting was unclear, a gene is shown on both sides. Irreversible steps of the glycolysis and gluconeogenesis are indicated in red.

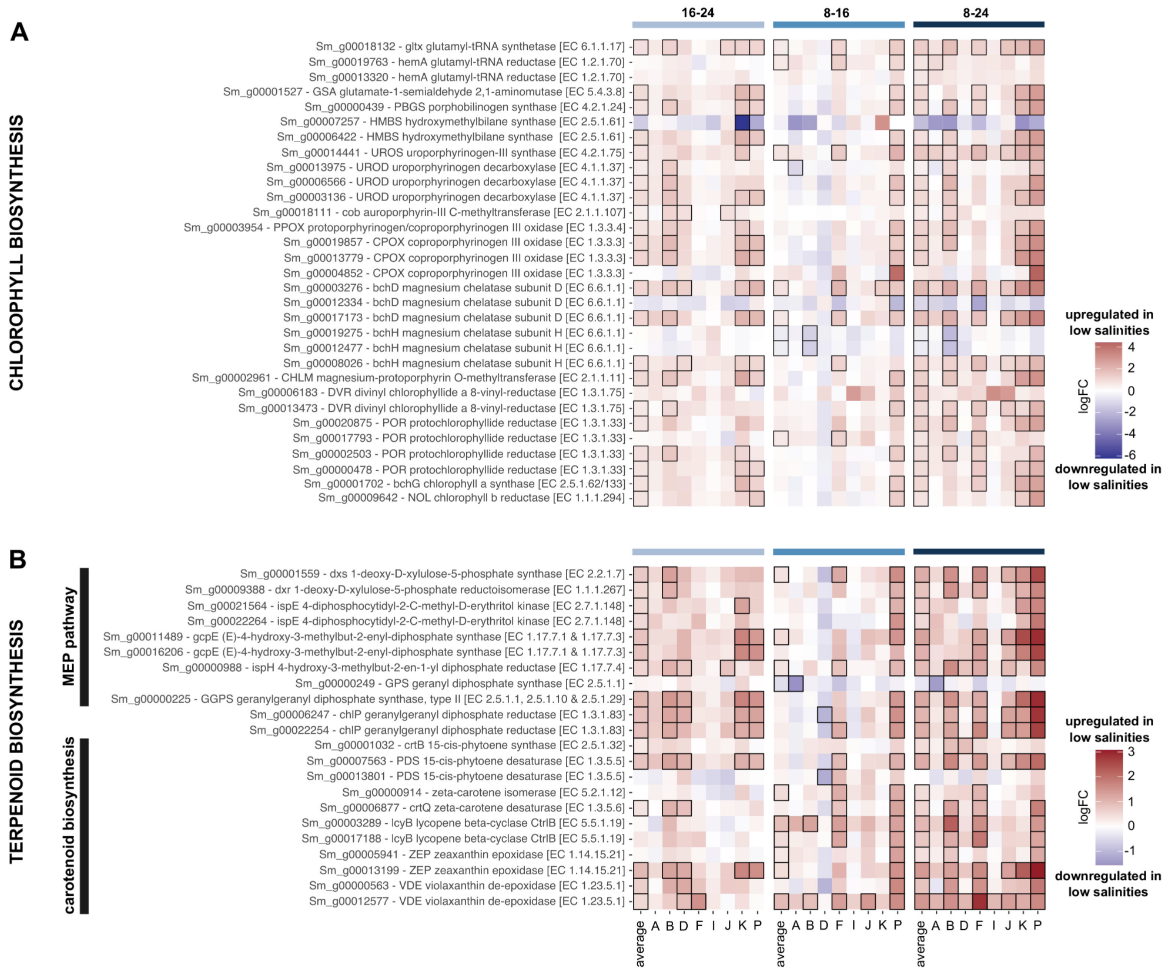


**Suppl. Fig. 4. Gene expression of genes involved in chlorophyll and terpenoid biosynthesis. a** Chlorophyll biosynthesis genes. **b** Terpenoid biosynthesis genes. Both the non-mevalonate (MEP) pathway and the downstream pathway to the biosynthesis of carotenoids are shown. The mevalonate pathway is not shown, as only a few genes of this pathway were DE. The heatmaps show logFC values of the 16-24, 8-16 and 8-24 contrasts of the average response and the eight genotypes. All visualized genes are DE in at least one contrast. Contrasts that were significant are outlined in black.

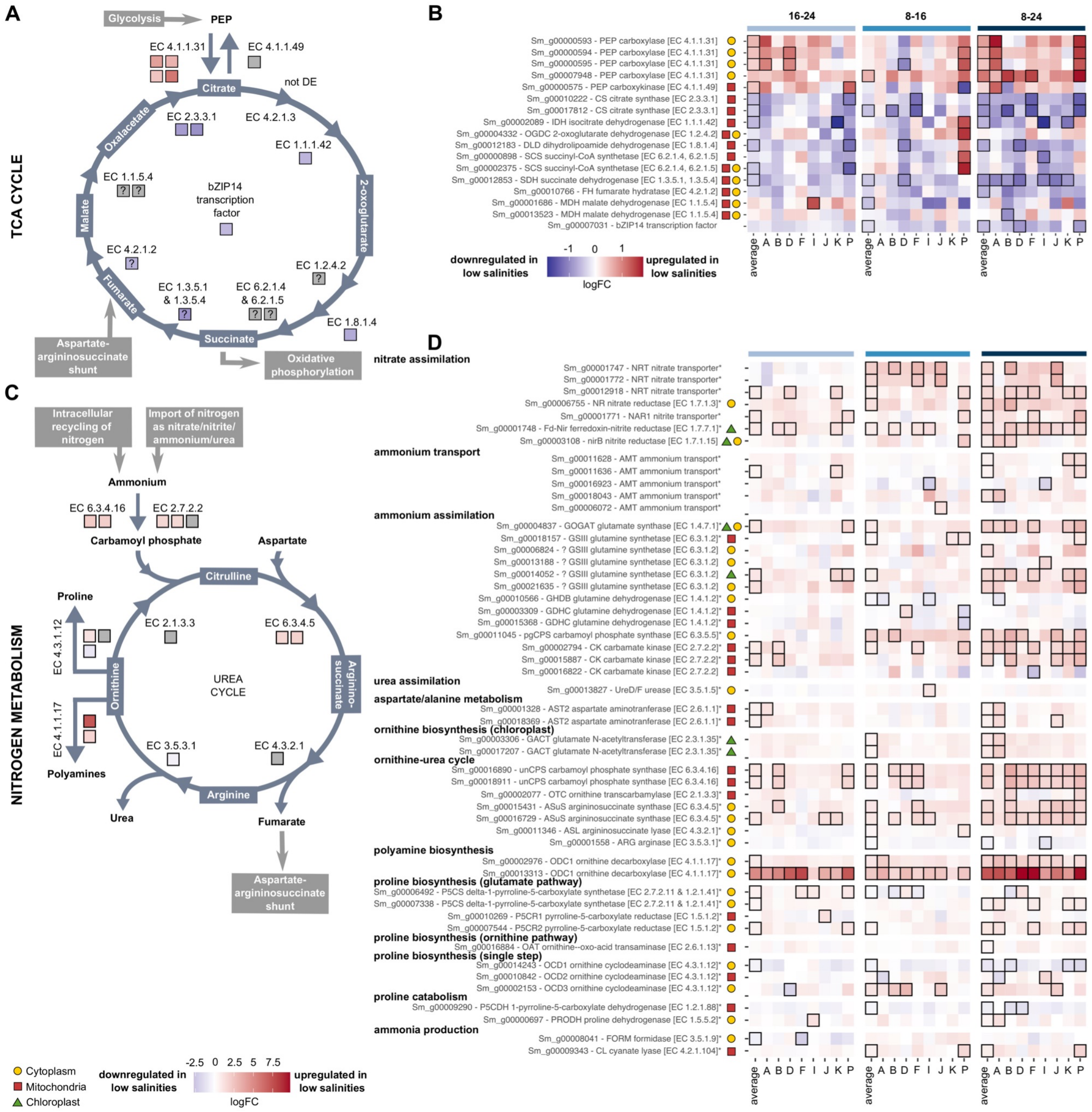

**Suppl. Fig. 5. Gene expression of genes involved in the TCA cycle and nitrogen metabolism, showing pathways and heatmaps on gene expression. a, b** TCA cycle. **c, d** Nitrogen metabolism, based on [37]. All heatmaps (**b, d**) show logFC values of the 16-24, 8-16 and 8-24 contrasts of the average response and the eight genotypes. All visualized genes are DE in at least one contrast. Contrasts that were significant are outlined in black. Symbols next to the gene names in the heatmaps denote protein targeting. In case the protein targeting was unclear, multiple symbols are used. Each colored square in the pathway figures (**a, c**) corresponds with a single gene, and is colored according to the logFC values of the 8-24 contrast of the average response. Squares filled with a question mark indicate proteins with unclear targeting.

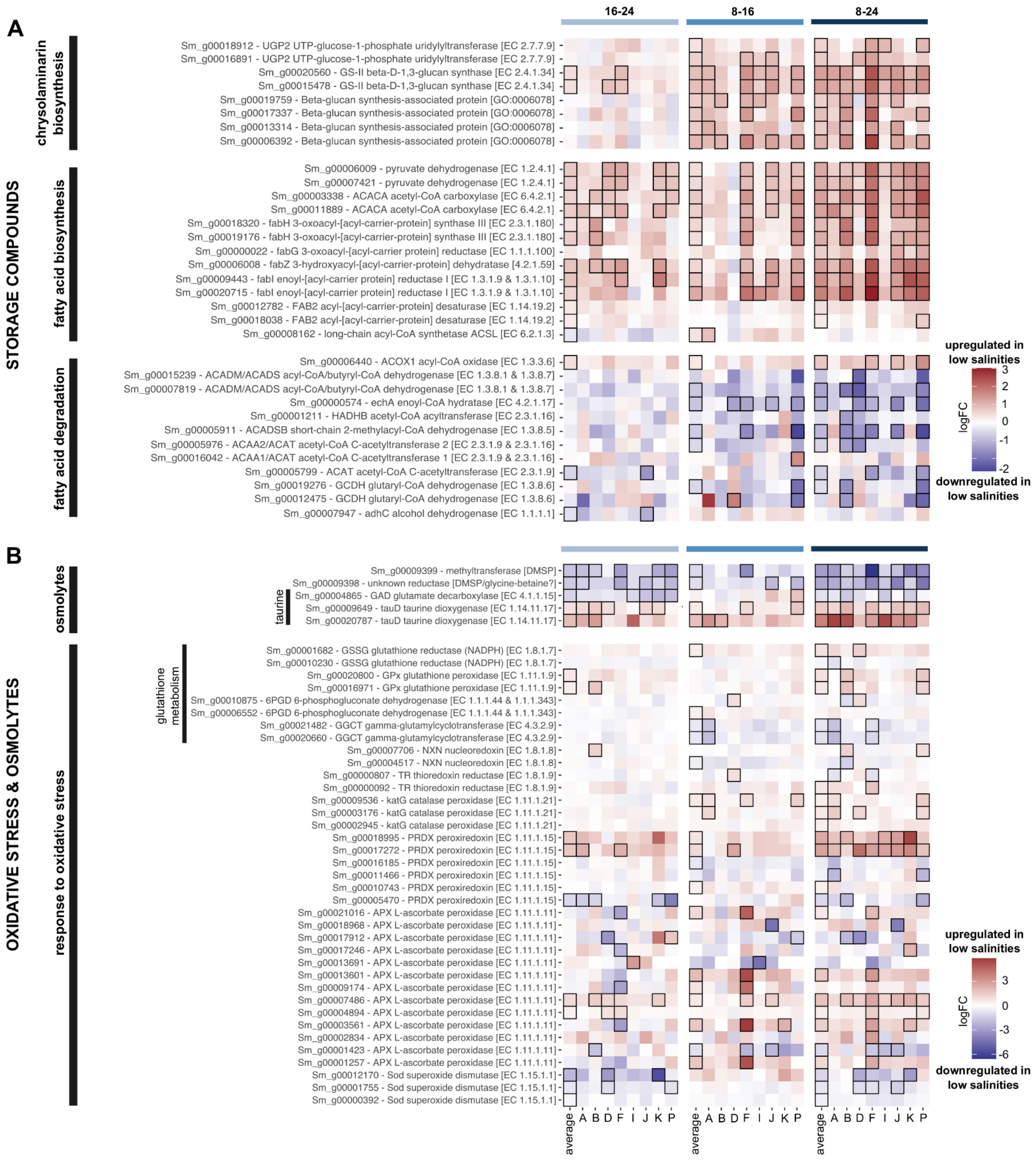


**Suppl. Fig. 6. Gene expression of genes involved in biosynthesis of storage compounds, osmolyte function, and response to oxidative stress.** **a** Biosynthesis of chrysolaminarin (β-1,3-glucans and β-1,6-glucans), and biosynthesis and degradation of fatty acids. **b** Osmolytes and oxidative stress. Proline and the xanthophyll cycle genes are not shown in (**b**), but are incorporated in the nitrogen metabolism (Suppl. Fig. 5) and terpenoid biosynthesis (Suppl. Fig. 4), respectively. The heatmaps show logFC values of the 16-24, 8-16 and 8-24 contrasts of the average response and the eight genotypes. All visualized genes are DE in at least one contrast. Contrasts that were significant are outlined in black.

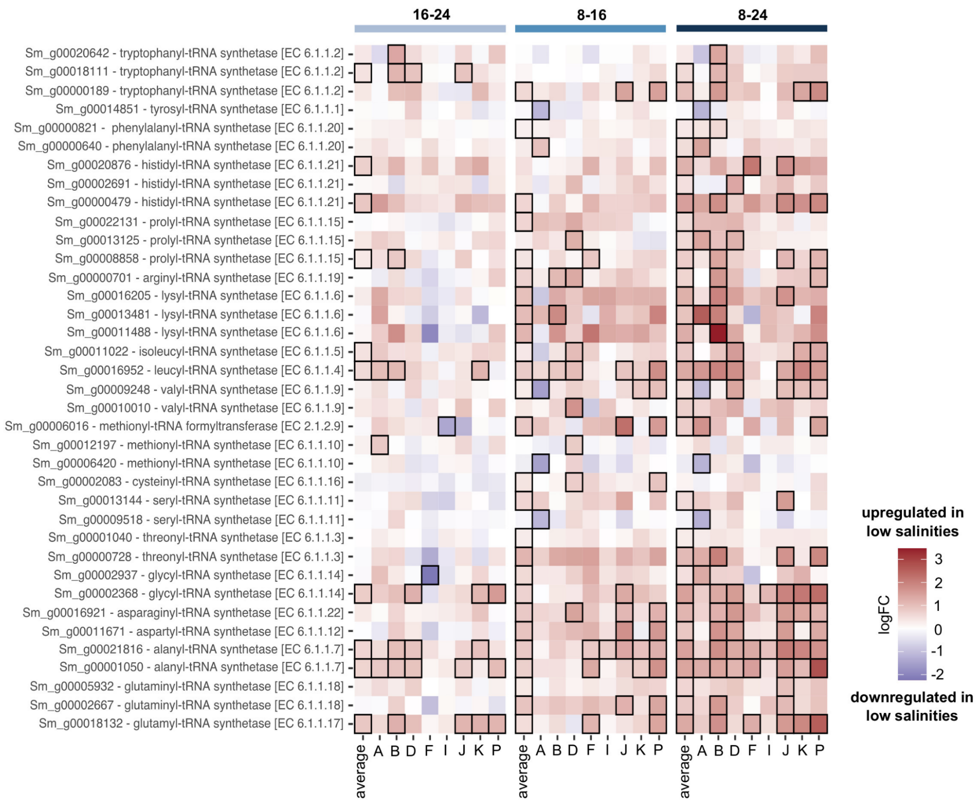


**Suppl. Fig. 7. Gene expression of genes involved in tRNA-aminoacylation.** The heatmap shows logFC values of the 16-24, 8-16 and 8-24 contrasts of the average response and the eight genotypes. All visualized genes are DE in at least one contrast. Contrasts that were significant are outlined in black.

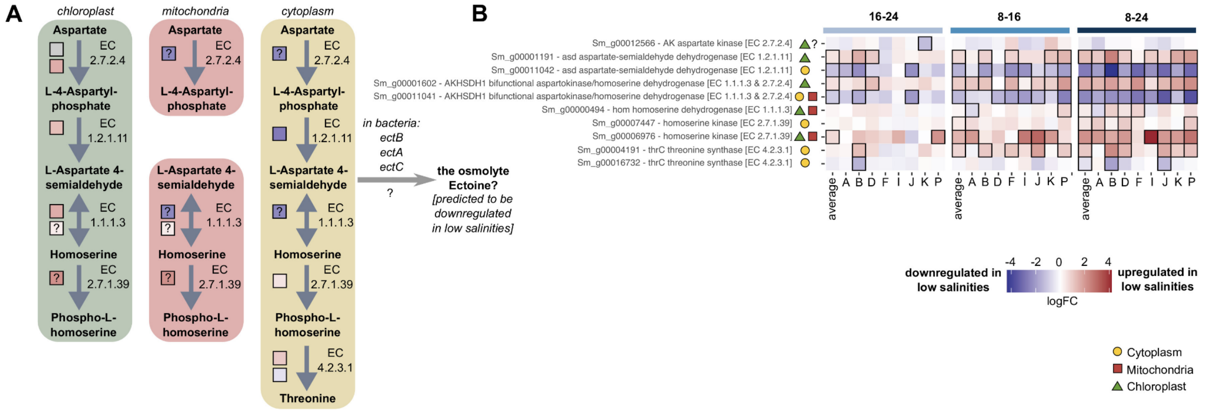


**Suppl. Fig. 8. Gene expression of genes involved in the first steps of the glycine-serine-threonine metabolism. a** Visualization of the first steps of the pathway, with distinction between the chloroplast, mitochondria and cytoplasm. **b** Heatmaps showing logFC values of the 16-24, 8-16 and 8-24 contrasts of the average response and the eight genotypes. All visualized genes are DE in at least one contrast. Contrasts that were significant are outlined in black. Symbols next to the gene names in the heatmaps denote protein targeting. In case the protein targeting was unclear, multiple symbols are used. Each colored square in the pathway figure (**a**) corresponds with a single gene, and is colored according to the logFC values of the 8-24 contrast of the average response. Squares filled with a question mark indicate proteins with unclear targeting. In case protein targeting was unclear, a gene is shown in multiple cell compartments. The grey arrow in (**a**) shows the presumed location of the branch towards ectoine biosynthesis.

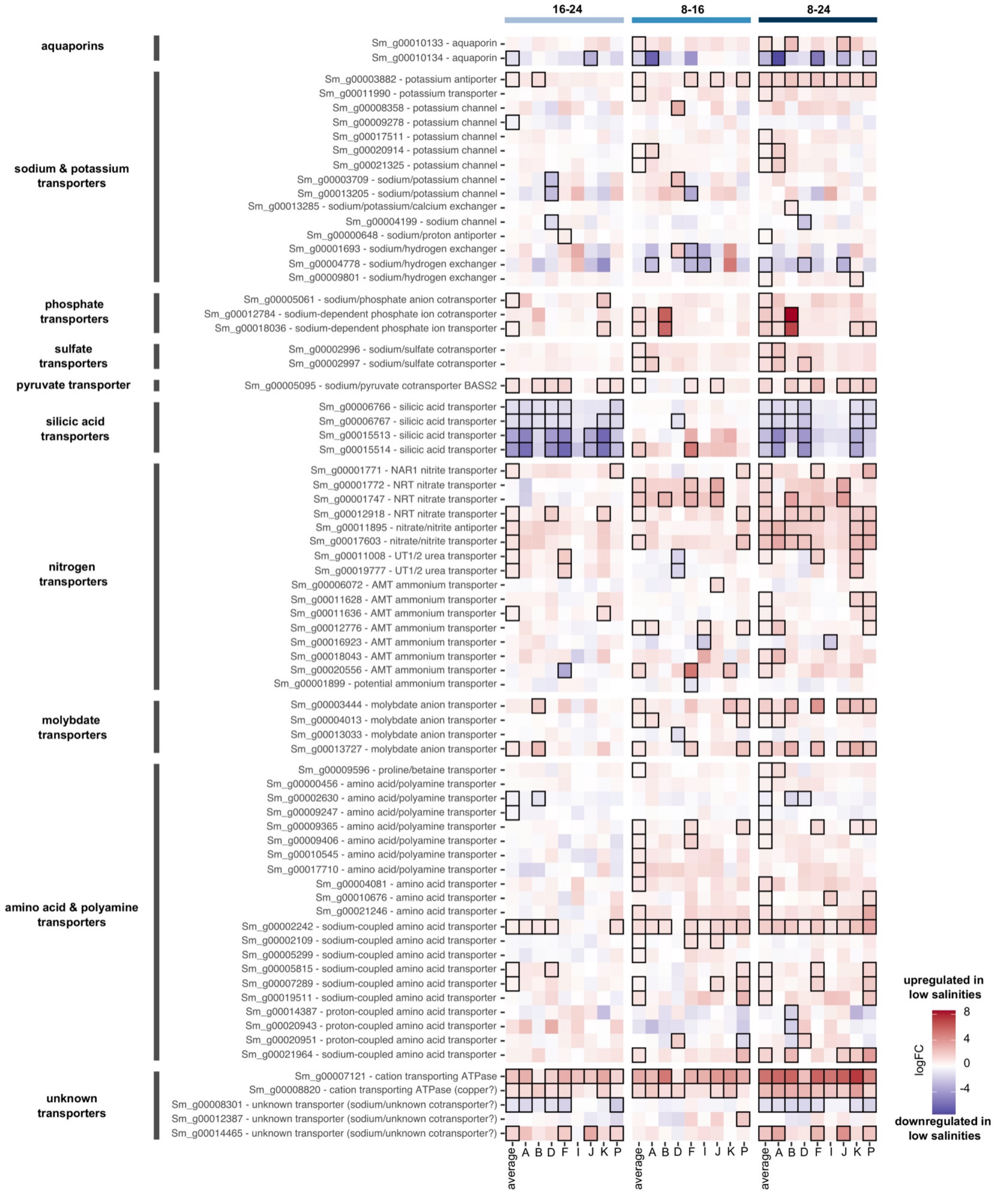


**Suppl. Fig. 9. Gene expression of transmembrane transporter genes.** The heatmap shows logFC values of the 16-24, 8-16 and 8-24 contrasts of the average response and the eight genotypes. All visualized genes are DE in at least one contrast. Contrasts that were significant are outlined in black.

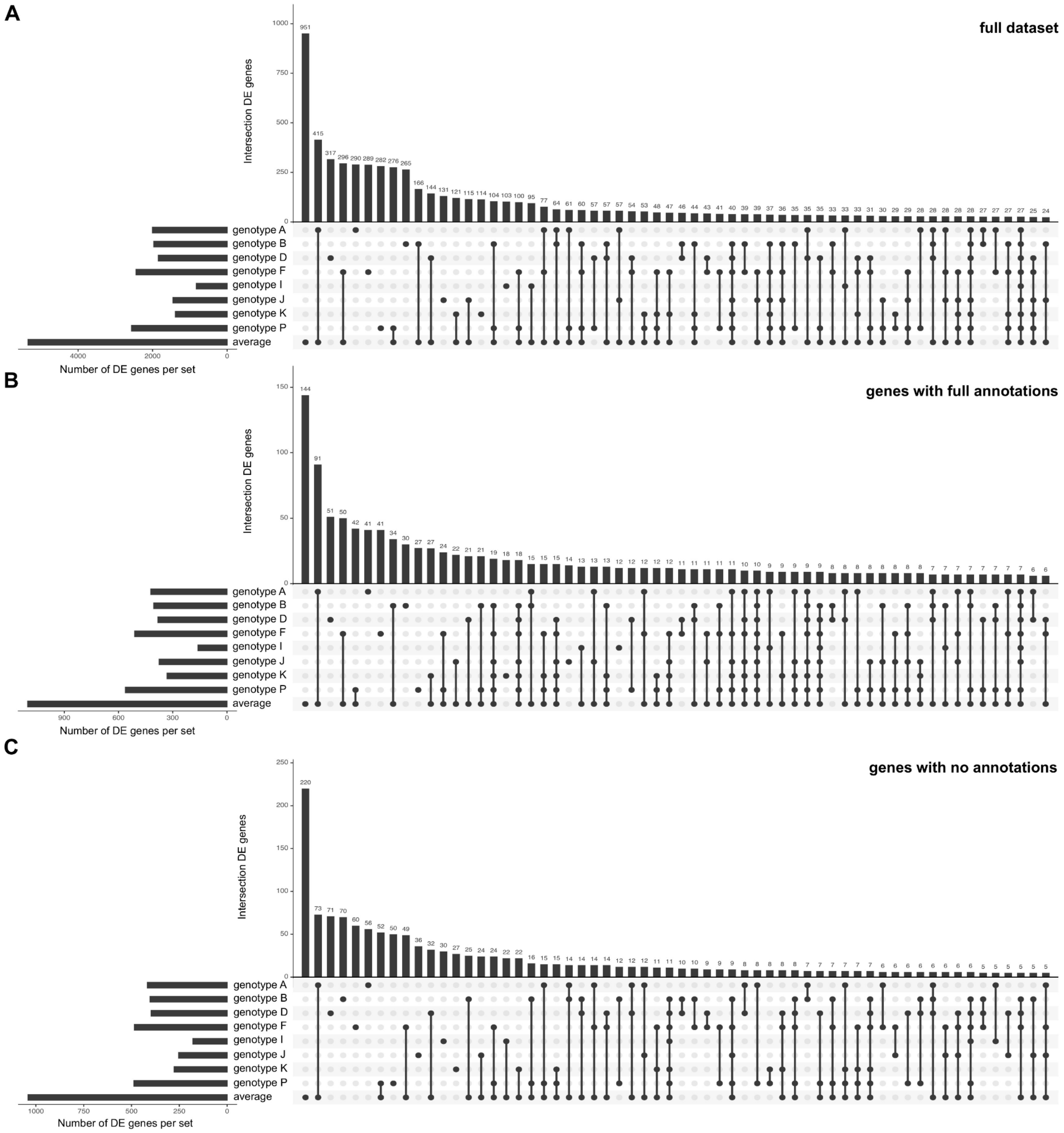


**Suppl. Fig. 10. Upset plots showing uniquely DE and shared DE genes between genotypes. a** Full dataset. **b** Genes with full annotation, i.e. with GO terms, KEGG annotations, Uniprot and Swissprot annotations, and at least one assignment to the InterPro, PANTHER, Pfam, SMART or SignalP databases as determined by InterProScan. **c** Genes with no annotations in the categories of set (**b**). For all upset plots, information from all salinity contrasts was combined.

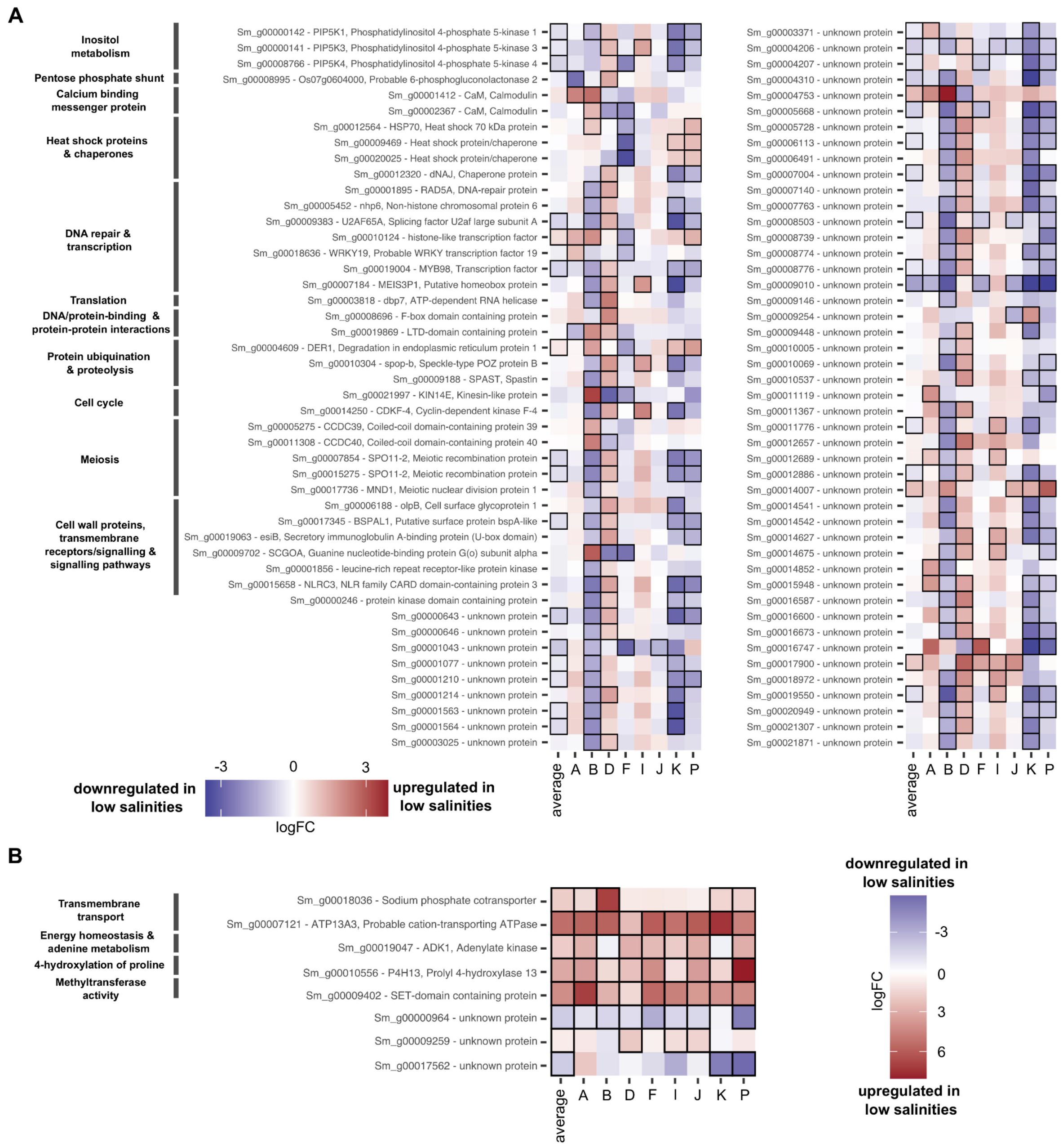


**Suppl. Fig. 11. Gene expression of the top 100 genes with significant interaction effects between genotypes.** Heatmap showing logFC values of the 8-24 contrasts of the average response and the eight genotypes. All visualized genes belong to the top 100 genes identified in the interaction-effect test, and were selected using stageR's FDR-adjusted P-value of the global null hypothesis (Padjscreen). When possible, functional annotations are indicated. Sixty-one of the top genes lacked (clear) functional annotation. **a** Genes which are DE in different directions in different genotypes. This set includes 92 genes. Note that some of the genes in the figure only show significant logFC values that are either positive or negative: these genes are classified as genes with DE in different directions because of significant logFC values in the 8-16 or 16-24 contrasts which are not shown in the figure (see Supplementary Methods for more information). **b** Genes which are DE in the same direction in different genotypes, but to a significantly different magnitude. This set includes 8 genes. Only contrasts significant in the genotype-specific responses were taken into account to classify genes into (**a**) or (**b**). Contrasts that were significant are outlined in black.

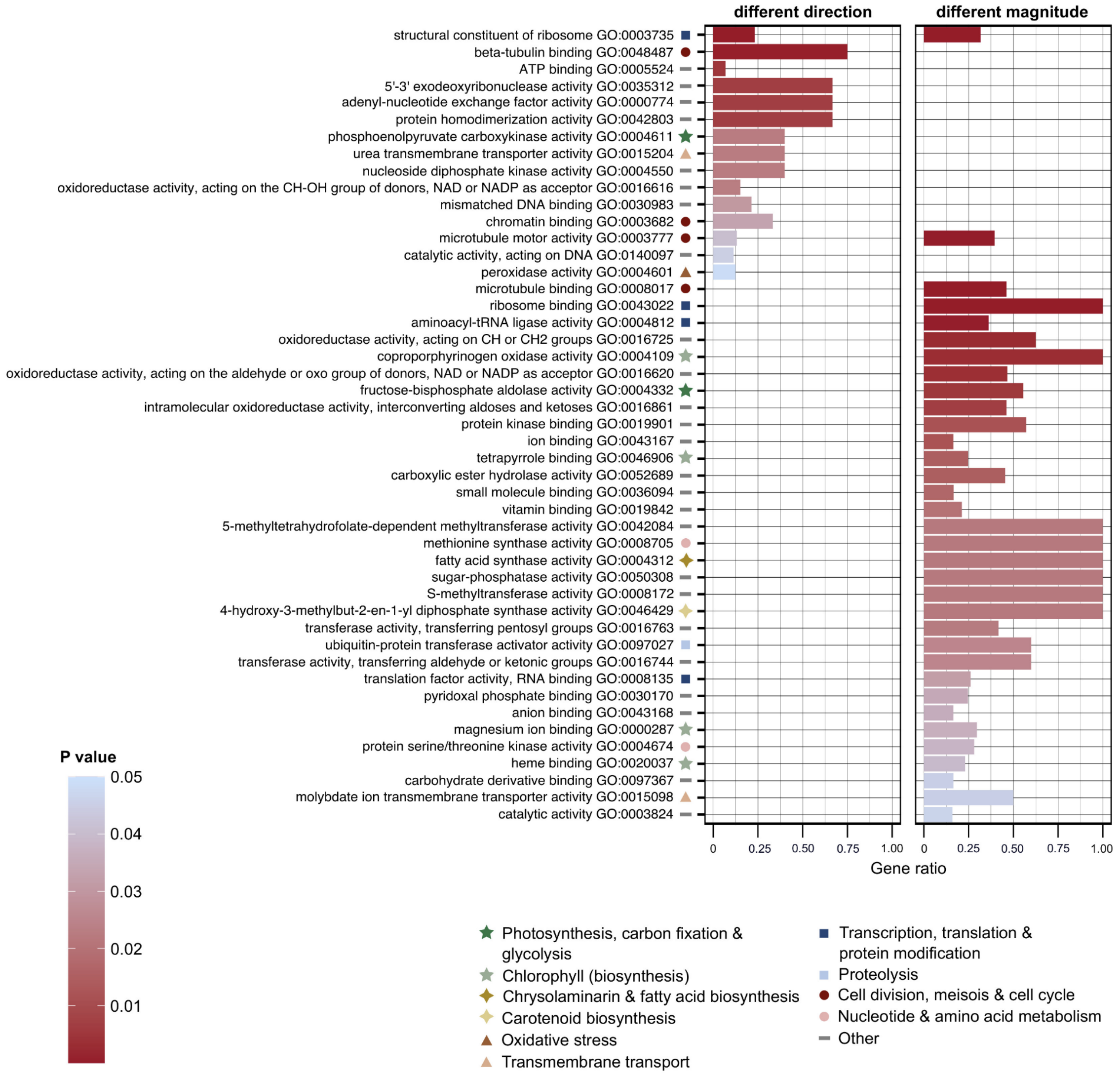


**Suppl. Fig. 12. GO enrichment on the interaction-effects: Molecular Function.** The barplots visualizes the significant GO terms retrieved by ORA (topGO, Fisher's exact test, *elim* algorithm) after removal of redundant GO terms by REVIGO. Two sets of GO enrichment were carried out which distinguished between genes that differ significantly between genotypes in the direction or magnitude of their response to low salinities. Barplot height indicates the proportion of genes that are DE with a given GO-term to the total number of genes with this GO-term in the genome of *S. marinoi*. The barplots are colored, and the GO terms ranked, according to P-value. Symbols indicate major categories of cellular processes to which a GO-term belongs. Only Molecular Function GO-terms are shown.

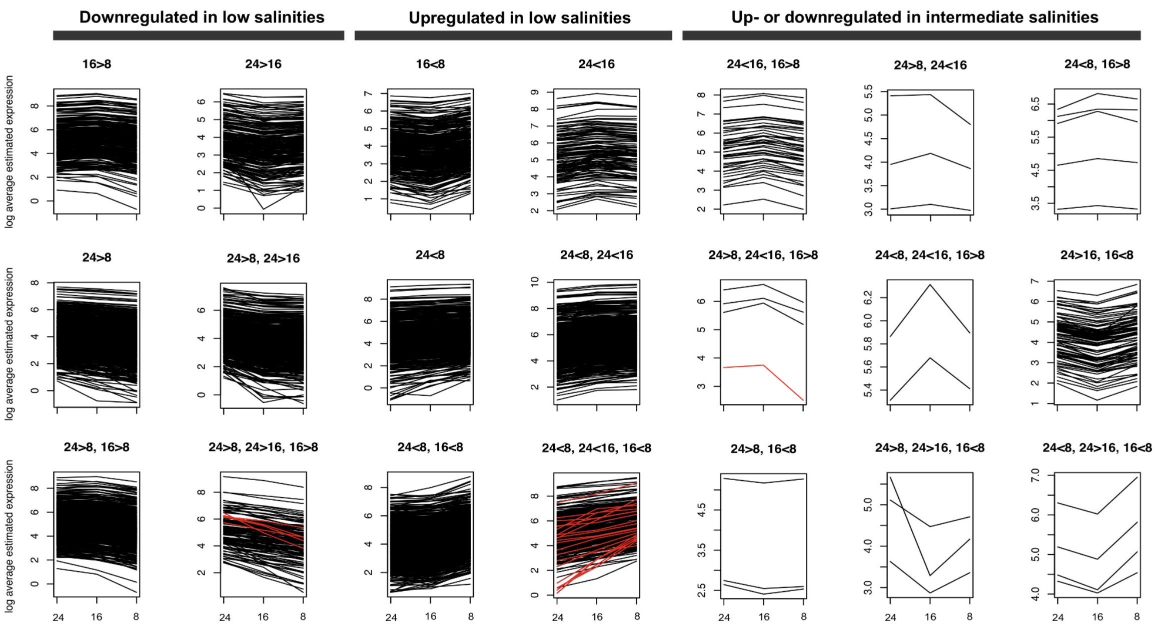


**Suppl. Fig. 13. TMM normalized logarithm of the average expression of each DE gene in function of salinity.** Gene expression data were averaged over all eight genotypes in this study using the log-fitted values of the glmQLFit output. Genes are assigned to different plots based on the DE contrasts and the direction of their response (e.g., genes in the 16>8 category are only DE in the 8-16 contrast and are always downregulated in salinity 8 relative to 16). Plots are ranked based on the overall direction of their response: downregulated in low salinities, upregulated in low salinities, and up- or downregulated in intermediate salinities. These first two sets were used as input for the ORA (in topGO) analysis on GO enrichment. Expression-curves indicated in red represent the 27 core response genes.
